# Current maladaptation increases with age in valley oak (*Quercus lobata*): Implications for future populations

**DOI:** 10.1101/2025.07.08.663605

**Authors:** Alexander R. B. Goetz, Marissa E. Ochoa, Jessica W. Wright, Victoria L. Sork

## Abstract

Adaptation and maladaptation are two fundamental evolutionary processes shaping the success of tree populations. Local adaptation has been studied extensively, but maladaptation, which not simply be the inverse, is particularly important for long-lived species since human impact is changing the environment on a very short time scale. Because maladaptation may be observed at different stages of an organism’s life cycle, temporal variation may reduce or enhance the vulnerability of a species to a new climate environment. In this paper, we use findings from an ongoing range-wide provenance study comprising two common gardens of 3,674 half-sib juvenile valley oaks (*Quercus lobata*) from 658 maternal families to address two objectives. First, we test whether maladaptation (a) persists as trees age and/or (b) varies in magnitude over time. We model growth rates and performance measures (height * survival; “performance”) across families sourced from sites with differing climates. Second, we assess the extent to which evidence-based seed transfer can mitigate the impact of maladaptation. We find that the twelve-year-old trees are currently adapted to temperatures not only cooler than their home site, but also beyond the coolest limit of the current species range. Moreover, this maladaptive pattern is exacerbated in hotter years of our study. Despite a general pattern of maladaptation, we demonstrate the feasibility of using phenotype-informed seed transfer guidelines for replanting trees in current populations by identified high-performing individuals from varied environmental origins. Overall, our results indicate that valley oak is imperiled by current and future climate conditions, but high-performing trees well suited to sites across the species range can be identified as seed donors. We conclude that current maladaptation in a long-lived tree species is a major threat that is first apparent within a few years and worsens as trees age, but careful management of vulnerable oak populations can improve their resiliency.

## Introduction

The increasing severity of extreme weather events and natural disasters, such as the recent wildfires in Southern California, presents an urgent danger to natural ecosystems of today as well as the future. Conservation projections and strategies often assume that organisms are optimally adapted to the environments in which they are currently found. In fact, thousands of studies have examined ways that selection has driven populations to become more fit to their environment, but the converse process—maladaptation—has been much less studied despite its impact on both evolutionary and ecological processes (Brady et al. 2019a). In addition, maladaptation is projected to affect many species in the future (Carter 1996, Davis et al. 2005, Aitken et al. 2008, Thuiller et al. 2008, Gougherty et al. 2021, Rellstab 2021, Sharma et al. 2022). Furthermore, mounting evidence has demonstrated that many species, including trees, are already maladapted to current climates (Rehfeldt et al. 2001, Savolainen et al. 2004, St Clair and Howe 2007). Given the variation in selective pressures on morphological and physiological traits as trees age (Barton 2024), combined with weather fluctuations across years (Wang et al. 2017), the extent of maladaptation may increase or decrease over time.

Quantifying the fluctuations in maladaptation is valuable to understand the long-term impact of tree response to local climate conditions and to apply evolutionary information to ecological conservation strategies. Maladaptation of tree populations to their environment poses a critical risk to species and ecosystem health. Since the classical experiments of Clausen, Keck and Hiesey (1940, 1947, 1948), many studies have documented local adaptation in plants (Sork 2018), but not all plants become locally adapted (e.g., Garrido et al. 2012, Etterson et al. 2020, Fréjaville et al. 2020, Gougherty et al. 2021), including tree species (e.g., St. Clair and Howe 2007). In tree species, maladaptation can result from historical adaptational lag (changes in climate outpacing evolutionary adaptation) causing genotypes to be distanced from their optimal climates (Davis et al. 2005, Hendry and Gonzalez 2008, Alberto et al. 2013). Long-lived tree species are especially likely to exhibit maladaptive patterns due to long generation times constraining genetic recombination (e.g., Frank et al. 2017, Browne et al. 2019, Gougherty et al. 2021, Rellstab 2021, Benomar et al. 2022, Schmeddes et al. 2024). Existing maladaptation, compounded by warming temperatures and habitat fragmentation, makes it even less likely that trees will be able to outpace altered climate regimes through either adaptation in place or natural gene flow (Davis et al. 2005, Aitken et al. 2008). Conservation strategies often assume local adaptation or uniform species-wide response to stressors (Benito Garzón et al. 2011, Breed et al. 2013). A critical question for conservation managers is whether populations are maladapted to current or future climates and whether such maladaptation affects future population growth, which may or may not be tied to the extent of evolutionary maladaptation (Brady et al. 2019b). Ideally, any efforts to restore or enhance tree populations will utilize climate-adapted genotypes that will both reduce evolutionary maladaptation in the population and increase population growth.

One additional question is important to address when a species is genetically maladapted to its environment: Can individual phenotypic plasticity “pick up the slack?” Plasticity in combination or opposition to genotype is a key aspect of a tree’s interaction with its environment, and can override the effect of underlying adaptive genetic variation on the phenotype (Chambel et al. 2005, Gimeno et al. 2008, Hamann et al. 2017). If plants can adjust to altered climatic conditions, they may show resilience despite genetic maladaptation. Indeed, genetic maladaptation cannot occur without sub-optimally fit genotypes persisting in their environment (Brady et al. 2019a). In addition, a high degree of phenotypic plasticity may mask the effects of underlying genetic maladaptation (Hamann et al. 2017). Plasticity in response to interannual climatic variation and/or longer-term acclimation (plastic responses over time leading to a better match to the local climate within an organism’s lifetime) to a planting site present possible “escape routes” from maladaptive patterns. Interannual fitness variation may result from differences in ontogeny (developmental changes throughout an organism’s lifetime; Quero et al. 2008, Barton 2024), which may in turn be either mediated or exacerbated by interannual climate variation (Xu et al. 2022). One way to assess the effects of plasticity is to document the extent to which plants respond to interannual climate fluctuations. If trees are highly plastic, we will not see declines in growth when short-term conditions differ from climates to which the trees are adapted (Alpert and Simms 2002). Conversely, vulnerability of trees to climate fluctuations would indicate that trees are “at the mercy” of changing abiotic environmental conditions. Therefore, it is possible that interannual variation in fitness can mitigate maladaptation to some degree if performance in “good years” is sufficient to bolster overall fitness and decrease the effect of “bad years.”

A constraint in quantifying maladaptation is that estimates can be biased if measurements happen at an insufficient temporal scale to capture background variation in growth (Bowman et al. 2013). Relationships between performance and origin climate at early tree life stages may be highly reflective of long-term patterns (e.g., Kashian and Barnes 2021, Chmura and Modrzyński 2023). Conversely, ontogenetic changes may greatly alter patterns as trees age (e.g., Germino et al. 2019), necessitating detailed longitudinal studies to disentangle potential sources of temporal variation. Furthermore, patterns of adaptation may be masked in the short term by maternal effects on progeny (Roach and Wulff 1987) or planting shock (Close et al. 2005), and plants may additionally acclimate to their planting sites over longer periods of time through plasticity (Sultan 1995). Despite these factors, genetic (mal)adaptation is typically quantified only once and often using young individuals (e.g., Gellie et al. 2016, Browne et al. 2019, Butnor et al. 2019, Bisbing et al. 2021). To our knowledge, no studies to date have explicitly quantified shifts over time in the extent of a species’ maladaptation to climate (but see e.g. Germino et al. 2019, Kashian and Barnes 2021 for studies of local adaptation in older individuals). Tracking interannual variation in growth and survival among trees will allow us to test whether individual plasticity can lead to improved or worsened outcomes in genetically maladapted populations.

Attempting to restore a foundational tree species without accounting for maladaptation is inherently risky amid increasing abiotic stress. Forestry has long recognized that locally adapted seed sources should be climatically matched to planting sites (Baldwin 1933), resulting in the common practice of using seed zones for replanting projects (Johnson et al. 2004). Given predicted warmer temperatures, recent work has called for instead selecting genotypes locally adapted to warmer parts of the species range for use in management and restoration of tree populations (Aitken and Whitlock 2013, Aitken and Bemmels 2016, Aitken et al. 2024). Using climate-informed approaches to seed sourcing can be a valuable tool for restoration projects (e.g. Breed et al. 2013, Gellie et al. 2016, Pedlar et al. 2021). Common gardens provide valuable information about seed sources for managing species of conservation and restoration concern (Schwinning et al. 2022). Empirically determining how seed source performance relates to climate can determine which genotypes are best adapted to future conditions and whether their degree of adaptation will be sufficient to promote population growth, thus ensuring that reforestation efforts are not hindered by unexpected maladaptation.

The overarching goals of this study are to test the hypothesis of maladaptation by measuring tree performance across years in a foundational California tree species, valley oak (*Quercus lobata* Née), and to test the hypothesis that sourcing genotypes from warmer climates can mitigate the species’ maladaptation to current and predicted future climates. Valley oak has already seen major reduction in its range due to land conversion (Kelly et al. 2005), and is predicted to undergo further contraction due to climatic shifts (Kueppers et al. 2005, Sork et al. 2010, McLaughlin and Zavaleta 2012). Here, we report results from a long-term provenance study of valley oak across two common gardens (Delfino Mix et al. 2015). Browne et al. (2019) found that these young trees were adapted to cooler temperatures, possibly due to lag-adaptation to historical climates, and predicted greater maladaptation in the future (Browne et al. 2019). Given that the trees were studied at only four years, it is possible that these growth patterns were a product of maternal effects (Roach and Wulff 1987), or that the trees have subsequently become acclimated to local conditions through plasticity (Sultan 1995). Thus, this study will investigate maladaptation using the same individuals in years 5-12 after germination and assess year-to-year fluctuations in growth and mortality associated with temperature conditions. Because of the large number of provenances represented in our two common gardens, this study provides a robust demonstration of intraspecific differentiation in adaptation to climate.

Specifically, this research utilizes empirical observations of growth and mortality of the half-sib progenies of 658 adult trees sampled from 95 localities measured during the first twelve years of the provenance trial to quantify temporal variation in maladaptation and test methods to mitigate its effects. We will explore four specific questions. **First,** are 12-year-old valley oaks adapted to current or previous temperatures? Valley oak will be considered adapted to current conditions if individuals perform best when the temperature difference between the maternal location and the common garden is close to zero. If valley oak is lag-adapted, we will extrapolate the curve of performance along the temperature distance gradient with a Gaussian function and determine the temperature where the curve is at its maximum. **Second,** does maladaptation decrease over time (suggesting acclimation) or are early signals of maladaptation indicative of long-term patterns? **Third,** does tree performance differ from year to year and does it decline during warmer years? To explicitly test performance of families from different environments, we will compare growth rates of cohorts of trees coming from warmer and cooler localities than the common gardens. **Fourth**, can performance of trees in a common garden be used to identify better-adapted genotypes for use in restoration? Our findings demonstrate that early warnings of species-scale maladaptation in a common garden are confirmed by long-term monitoring of performance, and that empirical growth data can improve conservation outcomes through phenotype-informed seed sourcing. The best evidence that maladaptation can be mitigated and result in future population growth are long-term studies that estimate survival and fecundity (Huxman et al. 2022). However, this study provides compelling initial evidence, based on tree growth as a component of fitness, that future population growth rates will be improved if restoration efforts utilize individuals with phenotypes that grow positively.

## Materials and methods

### Study species

Valley oak (*Quercus lobata* Née) is a long-lived (up to 600 years), winter-deciduous tree species endemic to woodlands, savannas, and riparian zones of the California Floristic Province, 0— 1700m above sea level (Pavlik et al. 1991). Though its historical range prior to European colonization is estimated to have covered much of the state, up to 95% of that range has already been lost to land conversion (Kelly et al. 2005) and only an estimated 3% of its remaining range is within protected areas (Davis et al. 2000). It is also known to be vulnerable to drought and increased temperatures (Kueppers et al. 2005, Tyler et al. 2006, McLaughlin and Zavaleta 2012), and shows evidence of local adaptation to drought (Mead et al. 2019). Furthermore, evidence shows that current valley oak populations are adapted to temperatures from the last glacial maximum rather than the recent past, but the degree of apparent maladaptation differs across genetic lines (Browne et al. 2019). Finally, shifts in local climatic niches of valley oak populations are predicted to outpace gene flow, limiting its ability to adapt (Sork et al. 2010).

### Common gardens

This study utilizes two common gardens established by JW Wright and VL Sork in 2012 as described in Delfino Mix et al (2015). In 2012, over 11,000 open-pollinated acorns were collected from 674 adult trees at 95 localities across the entire species range of valley oak (Figure 1, black dots). Acorns were germinated in 2013, grown for one year in a greenhouse at USDA Institute of Forest Genetics [IFG], Placerville, CA, (Figure 1, blue square) and transferred to lath houses for one year at IFG and the Chico Seed Orchard [CSO], Chico, CA (Figure 1, red square), and then planted into common gardens at CSO and IFG. Trees in the gardens are thus half-sib progeny of the maternal trees. CSO (70m above sea level) is hotter and dryer than IFG but has higher precipitation and lower winter temperatures than parts of the species range in southern California; IFG (822m above sea level) has a relatively cool and wet climate, but with hot summer temperatures and below freezing winter temperatures. In this paper, we refer to these as warmer (CSO) and cooler (IFG) gardens, respectively. Trees were planted according to a randomized block design to account for environmental differences between sections of each garden. Environmental differences among blocks (five per garden) are high, especially at the warmer site where two of the blocks are not contiguous with the others. Thus, our analyses account for these differences either statistically or through standardization (see below). Both gardens were irrigated from 2015 to 2019, and the warmer garden was irrigated throughout the entire study period. In 2021, roughly half of the individuals at each garden were thinned according to the original study design to prevent competition. As of 2024, the two gardens include a total of 3,674 surviving juvenile trees from 658 maternal lines.

**Figure 1:**
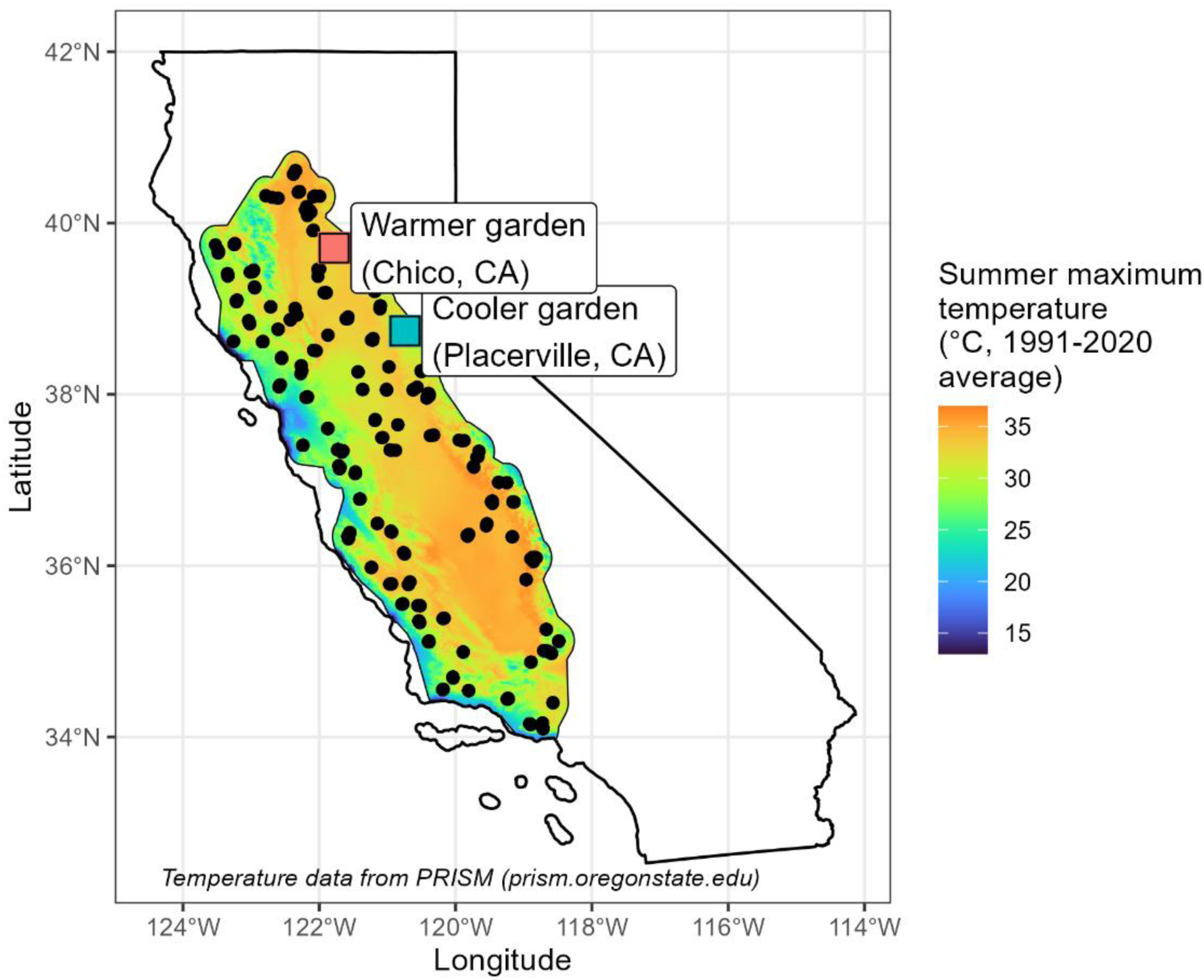
Maternal valley oak tree source locations (black circles; n = 95) within the current theoretical species range and common garden locations (red square: Chico Seed Orchard, Chico, CA; blue square: Institute of Forest Genetics, Placerville, CA). Background color represents 1991-2020 average of maximum yearly temperature (data from PRISM: prism.oregonstate.edu).

### Survival and growth data

At the end of the growing season in all study years, survival and size were measured (except for part of CSO in 2020 due to COVID-19 restrictions). Survival was assessed visually based on observable presence of living tissue; resprouts of individuals previously reported dead were common, so survival data were corrected afterwards. In the early years of the study, size was measured as height of the tallest stem; starting in 2019 (when trees were 7 years old), trees at least 140 cm tall had diameter at breast height (DBH) measured in addition to height. In 2021, trees removed for thinning had biomass destructively measured. These measures were used to estimate growth. Height was only measured up to a threshold in each year (Table S1), beyond which DBH was used to estimate height, with destructively measured biomass data used to supplement model training and validation (see below for details). See Table S1 for a detailed list of growth variables measured in each year. In 2018 at the warmer garden, DBH was only measured on a haphazard subset of trees, so we were unable to estimate height for those above the 4-meter threshold; we thus excluded the 2018 warmer-garden data to avoid biases. We also excluded all individuals that showed evidence of herbivore damage (especially a problem in the first year of growth in the gardens), as well as individuals that showed “negative” growth at any point due to either measurement error or dieback. Roughly 200 individuals out of 3,674 were excluded across all years as a result.

### Allometric growth calculations

To allow consistent comparison across different growth measurements, we fit allometric polynomial equations to the relationship between ln(basal diameter) and ln(height) of the trees destructively sampled in 2021 (Ochoa 2024), resulting in the following formula:

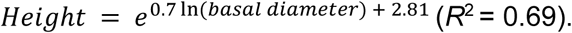

For comparison with other data, we converted DBH to basal diameter using an equation fit based on the relationship between 2018 basal diameter and DBH (basal diameter = 1.42(DBH) + 27.9; R^2^ = 0.72). We then used the basal diameter/height equation to predict height from DBH for trees lacking a true height measurement from 2018-2024. The R^2^ value of the relationship between predicted height and out-of-sample true height was 0.92. All subsequent calculations (growth rate and performance) were thus based on actual height when present (all individuals 2013-2017; individuals under the cutoff threshold in 2018-2024; individuals haphazardly sampled for complete height in 2023; all individuals at the cooler site in 2024) and height estimated from DBH otherwise. Partially due to inherent patterns of oak tree morphology, the models missed residual differences in the DBH/height relationship across individual trees. Predicted height also saturated at high DBH values. Finally, some trees had multiple dominant stems but only one stem per tree was measured (except in 2023), so predicted height may be underestimated in those cases. These constraints on our measurements of size and growth are unlikely to inflate the significance of our findings, so the trends we observe remain robust. (Refer to Table S1 for more details on growth variables across years.)

### Relative growth rates

To compare growth across trees from different lineages, we calculated relative growth rates (RGR) for each tree in the common gardens on both an annual and cumulative basis using the following equation:

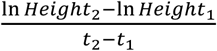

For the annual metric, t_1_ was the year before t_2_. If a tree had missing data for one year, the preceding year was treated as t_1_ (thus, t_2_ – t_1_ was always equal to either 1 or 2 in the interannual comparisons). For the cumulative metric, t_1_ was 2014 (the year before seedlings were outplanted in the gardens) and t_2_ was 2024. Prior to calculating RGR, we excluded any individuals with a recorded decrease in height or estimated height between subsequent years (generally caused by dieback or differences in rounding). We also excluded individuals that had died during the study period or that had been damaged by wildlife. Growth rates were not standardized by block, but differences among blocks were statistically accounted for in all tests by modeling a fixed effect of block nested within garden.

### Performance Indices: Multiplicative fitness function by block

To explicitly compare both growth and survival as fitness components of the maternal tree genotypes, we calculated multiplicative fitness functions (“performance”) for each family in each garden. Due to known differences in height across planting blocks, particularly at the warmer garden, we calculated individual standardized heights (dividing each tree’s height by the maximum height in its planting block, which in some cases was an estimated value) in each block and averaged across progeny of each maternal tree at each garden. Thus, the performance metric represents a component of the maternal genotype’s fitness at each garden. MFF was calculated using the following equations (*i* denotes individual progeny in gardens, *j* denotes maternal tree from which all *i* progeny descended).

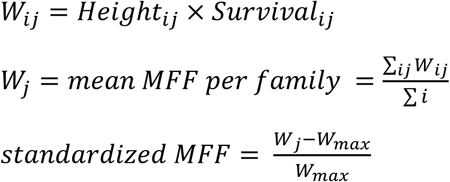

MFF was not calculated for the 2018 or 2020 data at the warmer garden since trees over 4m tall were not measured for height (DBH was only measured on a haphazard subset) and not all blocks could be measured in 2020 due to COVID-19 limitations.

### Temperature transfer distances

To compare the climates of the maternal seed sources to those of the gardens, we calculated transfer distances based on maximum summer temperatures and mean winter temperatures for each family in each year, taken as the difference in degrees between temperature in each garden each year and temperature in the 30-year averages of maternal source climate data. Maximum summer temperatures (among other climate variables) have been shown to constrain valley oak distribution (Kueppers et al. 2005, Sork et al. 2010, McLaughlin and Zavaleta 2012, Gugger et al. 2013) and are predicted to increase in the future (Wang et al. 2017). Mean winter temperature likewise reflects selective pressure imposed by freezing damage. We recognize that precipitation is also an important determinant of valley oak success (e.g., McLaughlin and Zavaleta 2012), but our analyses using precipitation showed the same trends (especially as precipitation and temperature were inversely correlated in the source sites). See Supplemental material, Figure S1 for precipitation trends in the gardens across the study period. Moreover, because the gardens were irrigated during the study period to ensure tree survival for analysis of traits and growth, we are using temperature variables. Finally, we also calculated summer temperature transfer distances based on temperatures averaged across June-August (see Supplemental material, Figures S2, S3), as well as correlations between summer and winter temperatures in the source sites (see Supplemental material, Figure S4).

Data on maternal tree source site climates from the California Basin Characterization Model (Flint et al. 2013) were used to compare common garden growth outcomes against historical climatic conditions experienced by the maternal source trees. All measurements were 30-year averages of 1950-1981 data gridded at 270m pixel resolution. Monthly climate data for the common garden sites were taken from PRISM (prism.oregonstate.edu). Paleoclimate data were taken from the CHELSA dataset (Karger et al. 2017). Projected future climate data (Relative Concentration Pathway 4.5; “stabilization” scenario) were taken from the AdaptWest ensemble projections, which use underlying data from the ClimateNA model (Wang et al. 2016, AdaptWest Project 2022). Note that here, unlike Browne et al. (2019), we present the temperature transfer distance in terms of the maternal source climate rather than the planting site (e.g., “trees from warmer climates than the garden” as opposed to “trees planted into cooler environments than the origin”) because it enhanced the clarity of the discussion of our findings. Regardless, the variable’s directionality and interpretation are the same across the two papers.

### Modeling effect of temperature difference on cumulative growth

To understand the effect of temperature transfer distance on lifetime growth of the trees in the gardens (Q1), we fit generalized additive models based on 2014-2024 growth rate and 2024 performance (as performance represents cumulative growth and survival up to the year of measurement). We used the same fitting methods as for the interannual variation models (see above). Models were fitted using the following formulas:

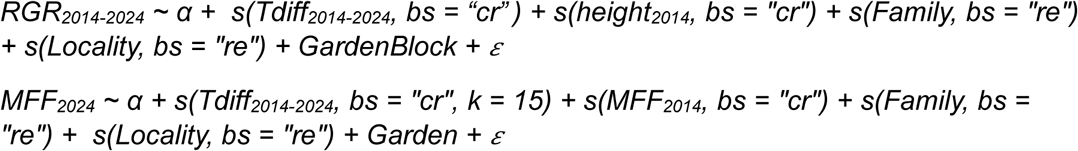

where *α* is the intercept and *ε* is the error term; RGR is growth rate; MFF is the multiplicative fitness function representing tree performance; Tdiff_2014-2024_ is temperature transfer distance between the source site average (summer maximum temperature, 1951-1980 mean) and each garden (summer maximum temperature, 2014-2024 mean).

### Modeling effect of temperature on inter-annual growth

To answer Q2 and Q3, we analyzed the effects of fluctuating maternal tree source site climates on growth rate using the generalized additive model framework (GAM; Hastie and Tibshirani 1986, Wood et al. 2015). As we were interested in examining the differences in climate responses across years, the independent variable of interest was the interaction between study year and temperature difference. Maternal tree ID and locality were included as random effects in the models to account for genetic and geographic differences between maternal trees. We also added block within garden as a fixed term to account for microclimatic differences within the gardens. We included initial height of the tree in 2014 as well as the height at the beginning of the interval used to calculate growth, to account for the fact that growth rate inherently decreases with increasing biomass (Rees et al. 2010). All numerical explanatory variables were standardized as Z-scores and modeled using splines with a cubic regression basis. We used Tweedie error distributions for model fitting due to the nonnormal shape of the response variable distribution (Browne et al 2019). The k value (knots used in fitting) was set to 15 for variables that indicated a poor fit with the default value (k = 10); the default was used for all other variables. We fit GAMs using the bam() function in package ‘mgcv’ (Wood 2011) in R 4.4.0/RStudio 2023.06.1 (RStudio Team 2023, R Core Team 2023). The model was fitted using the following formula:

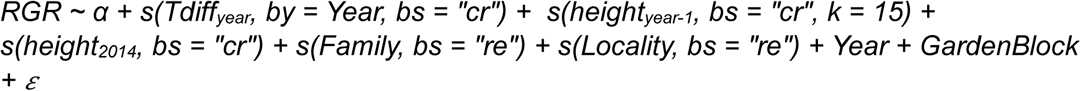

where *α* is the intercept and *ε* is the error term; Tdiff_year_ is temperature transfer distance within each year.

### Growth rate comparison across families and years

To determine whether differences in growth rates among families changed over time (Q2 and Q3), we conducted norm of reaction analyses that treated each study year as a separate environment. We conducted repeated-measures ANOVAs of the family*year interaction on individuals’ growth rates, controlling for planting block and with individual ID included as an error term, using the aov() function in R package stats.

To test whether hotter study years were associated with lower growth rates among families from different climate origins, we binned families into three transfer distance cohorts and compared their growth rates with annual garden maximum temperatures. Transfer distance values were calculated across all observations using the median summer maximum temperature within the study period (cooler garden: 2019, warmer garden: 2023). Cohorts were defined to represent discrete bins of transfer distance values, generally but not always in 5° C increments, to capture local modal transfer distance values as much as possible while also considering relative sample sizes (Figure S4). To validate the cohort groupings, we tested several series of similar cohort definitions, including random subsets of individuals. We consistently observed similar outputs, suggesting that the exact cohort definitions do not substantially influence the results. We then conducted regressions on the interaction between cohort and annual garden maximum temperature, controlling for planting block, using the lm() function in R package stats. Finally, we conducted the same regressions but excluding study years 3 and 4 (2015 and 2016) as all families exhibited very high growth rates in these years likely due to outplanting or ontogenetic effects. Gardens were analyzed separately since the cohorts were garden-specific.

To determine whether phenology influenced growth across years (Q3), we tested growth rate response to date of first leaf emergence, study year, and temperature transfer distance in each year. The model was fitted using data from both gardens, and included initial height and Block nested within Garden as fixed effects. The model also included source site locality as a random effect to account for regional differences; due to insufficient sample size of phenology data, we were unable to model differences among families.

### Extrapolating theoretical fitness maxima: Gaussian process regression

To aid in our interpretations of the cumulative and interannual analyses of the temperature/growth relationship, we used Gaussian process regression to extrapolate the conditions under which peak growth would be observed. Gaussian process regression is a nonparametric method that allows for extrapolation of models under the assumption that the modeled variable follows a normal distribution (Rasmussen and Williams 2006); Gaussian functions are used in quantitative genetics to mathematically describe the relationship between environment and fitness under local adaptation (Savolainen et al. 2007). To estimate theoretical fitness maxima in relation to the empirical relationship between fitness and temperature transfer distance, we used Gaussian process regression to extrapolate from the GAM fitted values of growth rate and performance. We fitted Gaussian process equations to each model (estimating mean and variance based on the shape of the GAM) and predicted values out to temperatures where the origin site would be 20°C warmer than the common gardens (the warmest origin sites were roughly 5°C warmer than the common gardens in the coolest study years). Gaussian process regression was fitted using function gpkm() with a Matern 5/2 kernel in R package ‘GauPro’, version 0.2.13 (Erickson 2024).

### Assessing seed sourcing scenarios

To answer Q4, we used data from the common gardens to simulate several strategies (random, climate-matching, and phenotypic selection) for planting seeds into a warm environment. First, we replicated the GAMs of cumulative performance in 2017, representing the 5^th^ year of the trees’ lifespans and the 3^rd^ year following outplanting in the gardens. Model was fitted using the following formula, where Tdiff_2014-2024_ = summer maximum garden temperature 2014-2024 – summer maximum source temperature 1951-1980:

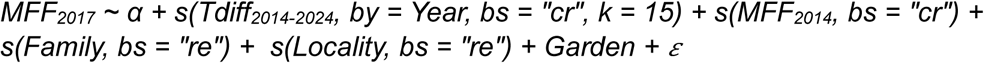

The random approach consisted of randomly selecting 100 families growing in the cooler garden. For the climate-matching approaches, we selected families with origin temperatures within ± 0.5°C of a) the cooler garden and b) the warmer garden. For the phenotypic selection approach, we selected the 90^th^ percentile of GAM-predicted performance in the cooler garden in 2017 (we also tested other quantile cutoffs and found similar results, with a general tradeoff between sample size and performance; see Supplementary material, Figure S5). We then tested differences in actual 2024 performance at the warmer garden among the seed source scenarios using a single-factor ANOVA and Tukey post-hoc test. We included seed source locality as a random error term in the ANOVA.

## Results

### Cumulative differences in growth and performance

To determine whether valley oak trees show evidence of maladaptation to temperature after ten years of growth (Q1), we modeled the effect of temperature transfer distance on cumulative growth rate and performance. We observed contrasting patterns of local adaptation to winter temperatures (Figure 2A, 2B) and maladaptation to summer temperatures (Figure 2C, 2D). Cumulative growth rate showed a clear pattern of peak growth in trees from origins with similar winter temperature to the gardens and progressively lower growth on either side of the peak (Figure 2A), while the performance metric (incorporating survival) showed an additional peak around +1°C winter T_diff_. Growth would theoretically be slightly lower under the predicted warmer future winter temperature increase of 1.3°C (Figure 2A, 2B, solid vertical lines). By contrast, when examining summer maximum temperature differences we observed faster growth rate (Figure 2C) and higher performance (Figure 2D) in trees from warmer source sites than the common gardens. The extrapolated relationships from Gaussian regression predict that the theoretical trait optima of growth rate and performance both occur closer to the summer transfer distance from the last glacial maximum temperature (−5.2°C) than to a transfer distance of zero (Figure 2C, 2D). Likewise, both trait values are much lower under a predicted future summer temperature increase of 1.8°C (Figure 2C, 2D). Analysis of average summer T_max_ showed comparable results (Figure S2). Full model summaries are provided in Tables S2-S7.

**Figure 2:**
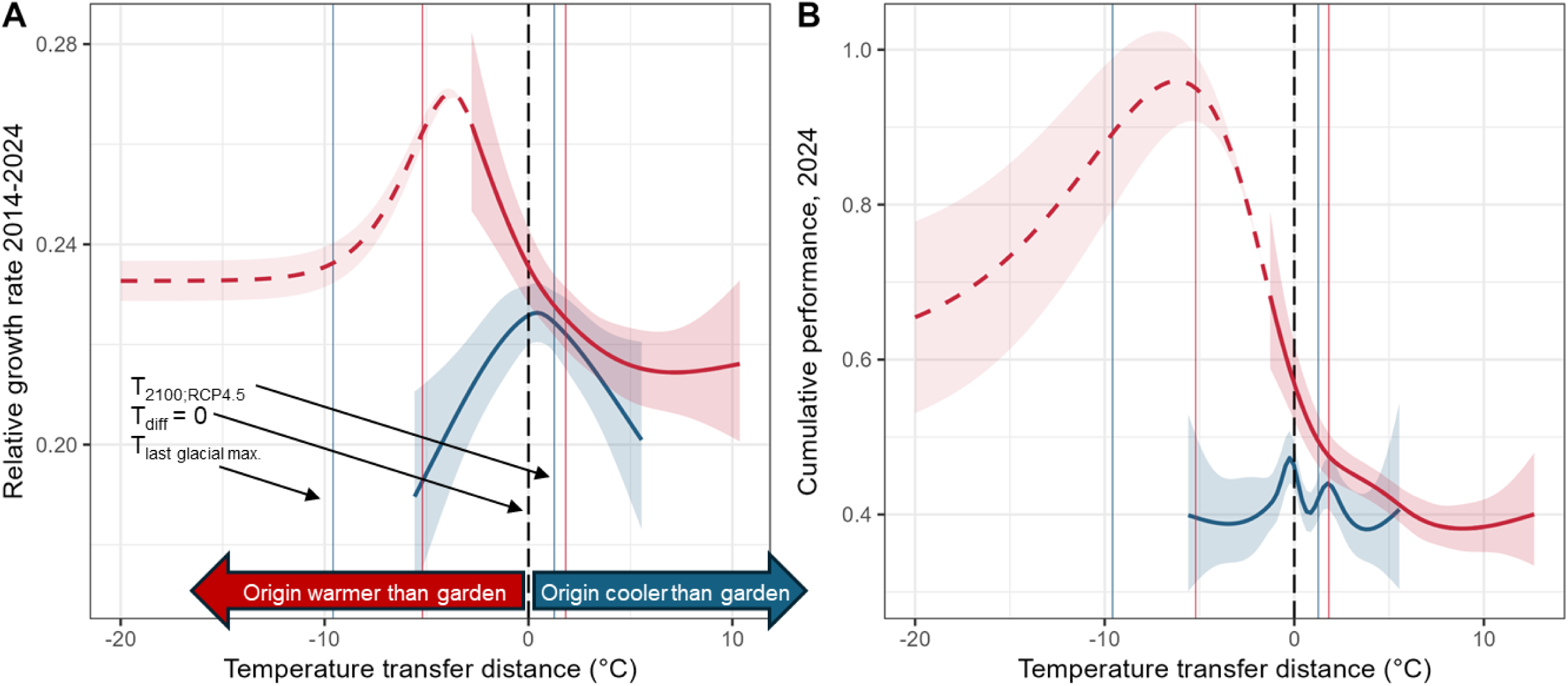
Ten-year cumulative trends in growth rate (left) and performance (right) as a function of temperature transfer distance (blue: mean temperature of coldest quarter; pink: maximum annual temperature) between maternal location and common garden. Solid curves denote generalized additive models fit to empirical data, and dashed curves denote extrapolation of the models using Gaussian process regression. Shaded regions represent 95% confidence intervals. The peak in each extrapolated curve represents the theoretical species-wide trait optimum. Vertical dashed line in each plot denotes where source temperature equals garden temperature; vertical solid lines denote estimated average temperature (blue: winter mean, pink: summer maximum) during the Last Glacial Maximum (21 kya; Wang et al. 2012) and predicted average temperature in year 2100 under a stabilization climate scenario (RCP 4.5; AdaptWest Project 2022). The area to the left of the zero line contains individuals from warmer historical climates than the garden, while the area to the right contains individuals from cooler historical climates. **A**: Relationship between mean temperature of coldest quarter and growth rate demonstrates local adaptation (maximum growth rate where origin temperature = garden temperature). Relationship between maximum temperature and growth rate shows maladaptation; extrapolated peak is close to the average temperature at the Last Glacial Maximum. **B**: Relationship between mean temperature of coldest quarter and performance shows peaks around both 0 transfer distance and temperatures slightly warmer than the origin. Relationship between maximum temperature and performance shows maladaptation; extrapolated peak is around 10°C cooler than the modern garden temperature.

### Interannual differences in growth rate and performance

To answer Q2 and Q3, we tested whether year-to-year variation in growth differs due to differences in climate each year. Winter mean temperatures significantly predicted growth rates in all but 2015, which had the warmest winter of the study period (Figure 3A). The significant relationships demonstrated a pattern of apparent local adaptation consistent with the cumulative analysis, in which the highest growth was observed in individuals with similar origin climates to the garden climates. However, while most years showed a local relative peak in growth around zero transfer distance, growth tended to be higher in families from warmer origins than the garden compared to families from cooler origins than the garden. Growth rates also differed significantly among years in relation to garden/source summer temperature differences (Figure 3B). For example, in one of the cooler years, 2016, families coming from cooler sites than the gardens show a relative peak in growth rate. However, in most years, the individuals from warmer sites than the common gardens had higher growth rates. Mean growth rate was also variable and tended to be higher in cooler years. The extrapolated relationships show an estimated peak in growth rates close to the temperature transfer distance from the last glacial maximum (−5.2°C) in all but the two coolest years. In most years, predicted growth rate was lower at transfer distance of +1.8°C (the degree of warming predicted by 2100 under the Relative Concentration Pathway 4.5 “stabilization” scenario) than 0°C transfer distance, but the difference was relatively small (e.g., roughly 1% in 2023). However, in 2021 (the second hottest year of the study to date) the predicted growth rate at +1.8°C transfer distance was roughly 23% lower than at 0°C transfer distance. Full model summaries are provided in Tables S8, S9. The effects of June-August mean transfer distance on growth rate were very similar to the effects of overall maximum temperature transfer distance (Figure S3; Table S10).

**Figure 3:**
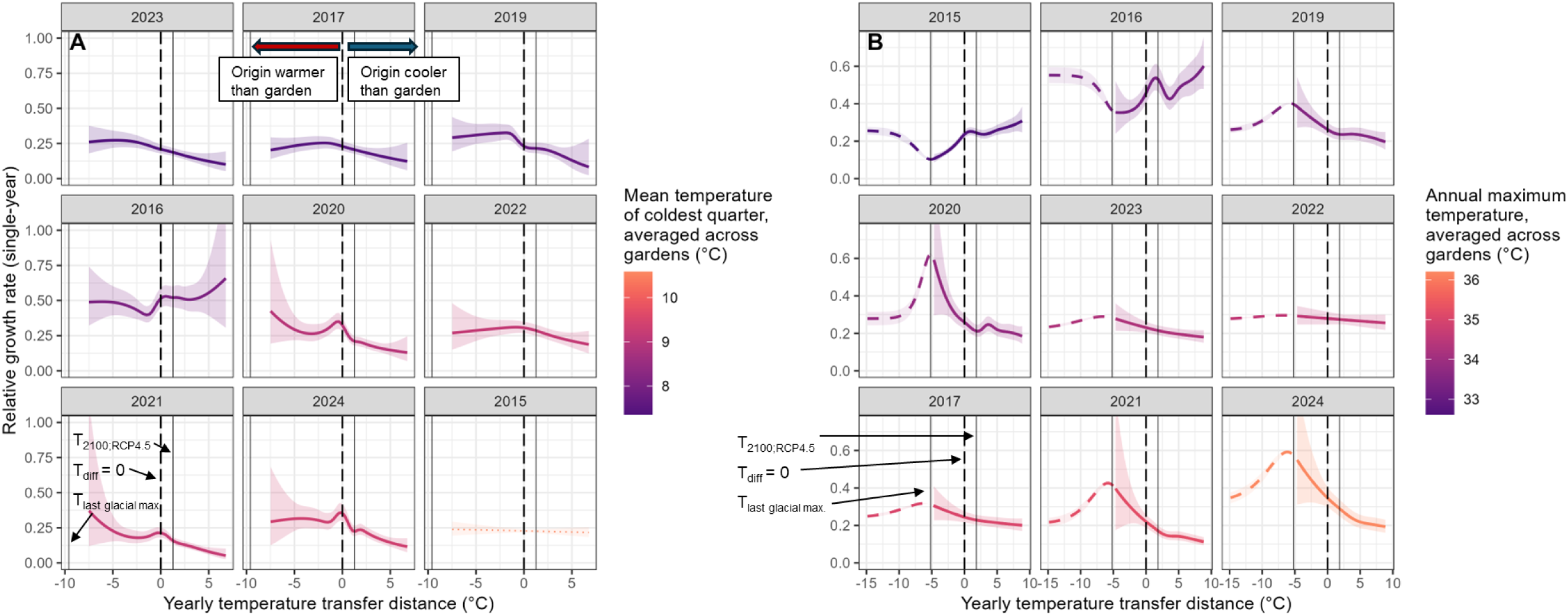
Yearly trends in valley oak growth rate as a function of transfer distance between maternal location and common garden, arranged and color-coded in order of cooler to warmer temperatures. Solid colored curves represent empirically derived generalized additive model predictions of single-year relative growth rate as a function of yearly temperature transfer distance (**A:** mean temperature of coldest quarter; **B:** maximum summer temperature) across the period of tree growth in the common gardens, 2015-2024 (2018 is excluded due to incomplete data). Long-dashed curves represent extrapolations of the model using Gaussian process regression. Thin dotted curves represent non-significant relationships. Vertical dashed line in each plot denotes where source temperature equals garden temperature; vertical solid lines denote estimated average temperature during the Last Glacial Maximum (21 kya; Wang et al. 2012, Karger et al. 2017) and predicted average temperature for years 2070-2100 under a stabilization climate scenario ensemble (RCP 4.5; AdaptWest Project 2022). The area to the left of the zero line contains individuals from warmer historical climates than the garden, while the area to the right contains individuals from cooler historical climates.

We observe anecdotally that the shapes of the predicted temperature/growth rate relationships are similar among years with similar mean precipitation, even if temperature was different. For example, mean precipitation across gardens in 2017 and 2019 was 1207mm and 1218mm respectively, but mean temperatures were very different (35.0°C vs. 33.5°C). Nonetheless, the shapes of the two years’ predicted relationships were more similar than those among the three hottest or three coolest years (Figure 3B). Likewise, despite being an “average” year in terms of temperature, 2020 was the driest year of the study (450 mm of precipitation), and had the sharpest predicted performance decline with increasing distance from the peak (greater penalty for non-optimally adapted genotypes). We point out that precipitation may be playing a role in common garden performance outcomes, but we report these findings consciously because the gardens were irrigated during the early years to ensure that trees would survive for purposes of the experiment. Nonetheless, we see this trend associated with the precipitation of the maternal site and present the pattern of 2014-2024 annual precipitation at the common gardens in Figure S1.

To directly test the genetic effect of family in relation to interannual variation on growth rate (Q2, Q3), we analyzed the reaction norms of growth rate over time, treating each year’s climate as a separate environment (Figure 4A, 4B). We found significant effects of year and family, but not year*family interaction, on growth rate at both gardens when controlling for planting block (Tables S11, S12). The statistical results indicate that growth rates changed among years and families differed in their responses to interannual variation, but their responses did not shift in relation to each other over time (i.e., each family tended to grow either slowly or quickly in all years relative to other families). We also see a large peak in growth rate across all families in 2016, likely due to ontogeny (fast growth at the second-year seedling stage) or acclimation to the site (2016 was the second growing season after out-planting). While we could not directly separate the effects of tree ontogeny and climate, higher annual maximum temperatures were significantly associated with lower growth, even when 2015 and 2016 were excluded from analysis (Figure 4B, 4C). Temperature transfer distance cohort was only weakly predictive of the relationship between growth rate and annual temperature; in general, differences among years were much higher than distances among cohorts (Figure 4B, 4C).

**Figure 4:**
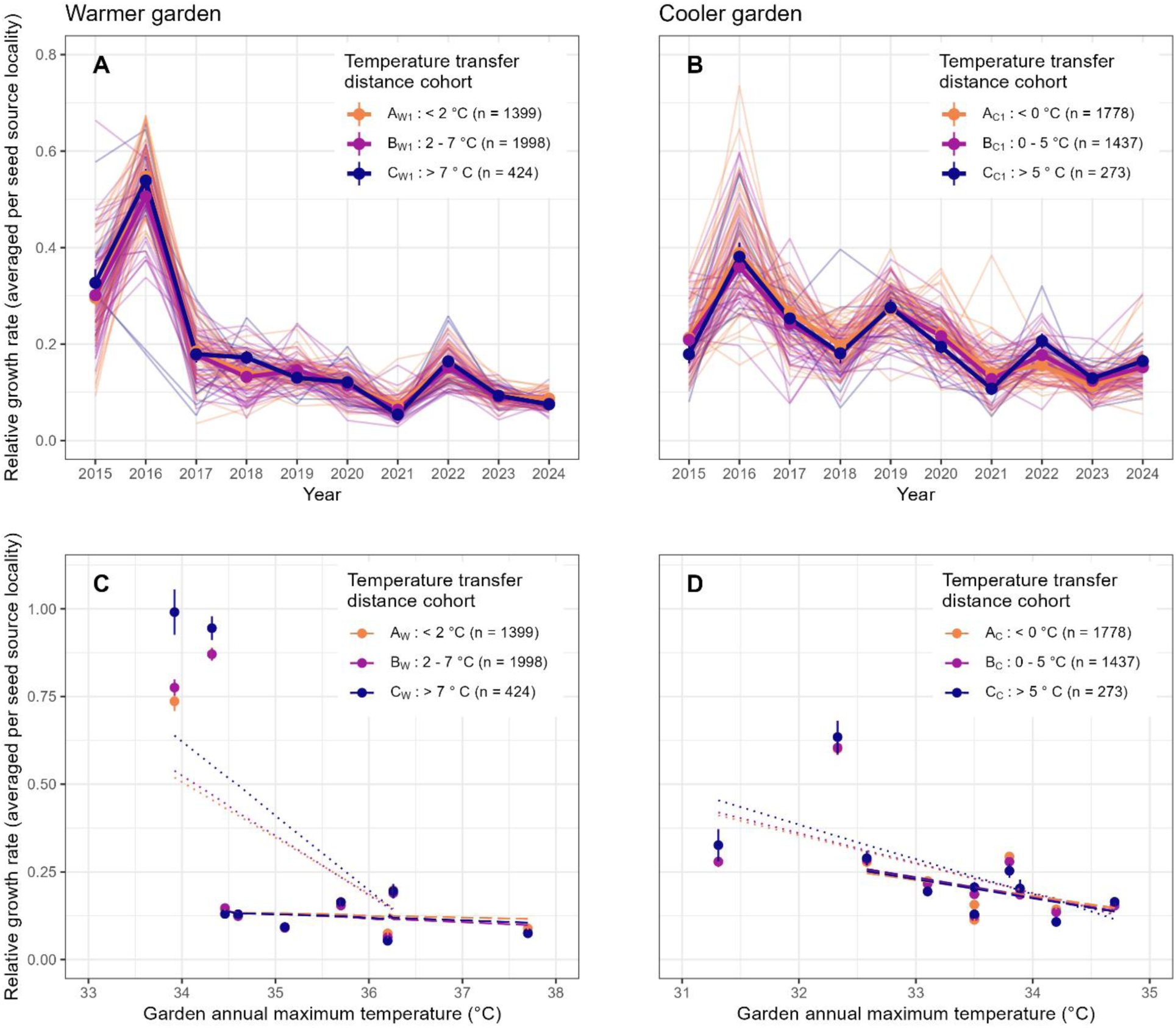
**A-B**: Norms of reaction across years for each garden showing average annual relative growth rate per year of each source site locality (thin colored lines). Line colors illustrate cohorts based on temperature transfer distance between garden and source site. Thick lines and points show cohort-wide means (± SE) of relative growth rate. **C-D:** Scatterplots with regression lines illustrating effect of garden annual maximum temperature on annual relative growth rate. Points show mean ± SE of growth rate for each cohort in each year. Regression lines are plotted for each cohort; regressions on all years are shown as dotted lines, while regressions excluding study years 1 and 2 (the years with the highest growth overall, likely due to planting effects) are shown as long-dashed lines.

To determine whether phenology predicted fitness outcomes (in particular, whether fitness differences were only due to differences in growing season duration; Q3), we tested the relationships between climatic transfer distance, phenology, and growth rate in each year. Date of first budburst differed across study years and by family and locality (Table 1). We observed that families from warmer climates had earlier leaf emergence than those from cooler climates. In addition, relative growth rates were higher in families with earlier leaf emergence across both gardens when controlling for climatic distance between source site and garden (Table 2). In addition, earlier leaf emergence date was associated with higher garden maximum temperatures during early spring when controlling for planting block within garden, year, and maternal seed source (February-April; *p* < 0.001, R^2^ marginal = 0.59, R^2^ conditional = 0.67). These findings indicate that length of growing season contributes to higher growth rates but is not the only mechanism that explains the difference between families of varying climatic origins.

**Table 1:** Date of first budburst differs among years and by family nested within locality (random effects). Estimates are estimated budburst date in each year (relative to 2015 budburst date estimate). σ^2^ = within-group variance; τ_00_ = between-group variance; ICC = intra-class correlation coefficient. Planting block, nested within garden, was included to account for unintended environmental differences within gardens but is not shown here for easier interpretation.

| Date of first budburst |  |  |  |
| --- | --- | --- | --- |
| *Predictors* | *Estimate* | *CI* | *p* |
| (Intercept) | 72.74 | 71.78 – 73.70 | <0.001 |
| Year [2016] | 2.44 | 1.55 – 3.33 | <0.001 |
| Year [2018] | 19.91 | 19.00 – 20.81 | <0.001 |
| Year [2019] | 16.14 | 15.36 – 16.92 | <0.001 |
| **Random Effects** |  |  |  |
| $\sigma^2$ | 91.69 | | |
| $T_{00}$ Family:Locality | 6.29 | | |
| $T_{00}$ Locality | 14.53 | | |
| ICC | 0.19 |  |  |
| $N_{\text{Family}}$ | 619 | | |
| $N_{\text{Locality}}$ | 95 | | |
| $N_{\text{observations}}$ | 3732 | | |
| Marginal $R^2$ / Conditional $R^2$ | 0.389 / 0.502 | | |

**Table 2:** Linear mixed model of valley oak trees’ individual relative growth rates as a function of year, temperature transfer distance, source site locality, and phenology. Estimates are relative growth rates estimated for each model term. σ^2^ = within-group variance; τ_00_ = between-group variance; ICC = intra-class correlation coefficient. Planting block, nested within garden, was included to account for unintended environmental differences within gardens but is not shown here for easier interpretation.

| *Predictors* | **Relative growth rate** |  |  |
| --- | --- | --- | --- |
|  | *Estimate* | *CI* | *p* |
| **(Intercept)** | 0.85 | 0.81 – 0.88 | **<0.001** |
| **Date of first budburst** | 0.02 | 0.00 – 0.03 | **0.007** |
| **Year [2016]** | 0.23 | 0.20 – 0.25 | **<0.001** |
| **Year [2018]** | -0.40 | -0.43 – -0.37 | **<0.001** |
| **Year [2019]** | -0.36 | -0.39 – -0.33 | **<0.001** |
| **Temperature maximum difference (garden – source)** | 0.02 | 0.01 – 0.03 | **0.001** |
| **Random Effects** |  |  |  |
| $\sigma^2$ | 0.15 | | |
| $T_{00}$ Locality | 0.00 | | |
| ICC | 0.01 |  |  |
| $N_{\text{Locality}}$ | 95 | | |
| $N_{\text{observations}}$ | 8787 | | |
| Marginal $R^2$ / Conditional $R^2$ | 0.332 / 0.336 | | |

### Hypothetical seed sourcing scenarios

To answer Q4, we compared fitness outcomes of a simulated cool-to-warm transplant in which performance of the same families were compared between year 5 at the cooler site (simulating a source population) and year 12 at the warmer site (simulating a planting site). Selecting families by matching origin climate to either garden climate resulted in performance higher than the overall mean but not significantly different from the random selection approach (Figure 5). Furthermore, there were no families with origin temperatures at least 2°C hotter than the 2014-2024 average temperature at the warmer site (i.e., no options for explicitly climate-adaptive seed transfer). By contrast, selecting families in the 90^th^ percentile for growth predicted by the T_max_ difference GAMs resulted in significantly higher than random selection (Figure 5). Furthermore, this category included several families from cooler origin sites than the gardens. Finally, as the high-performing families could be successfully identified in year 5 of the study, we find support for the use of short-term seedling studies to identify successful phenotypes with potential for use in reforestation. Full model summary is provided in Table S13.

**Figure 5:**
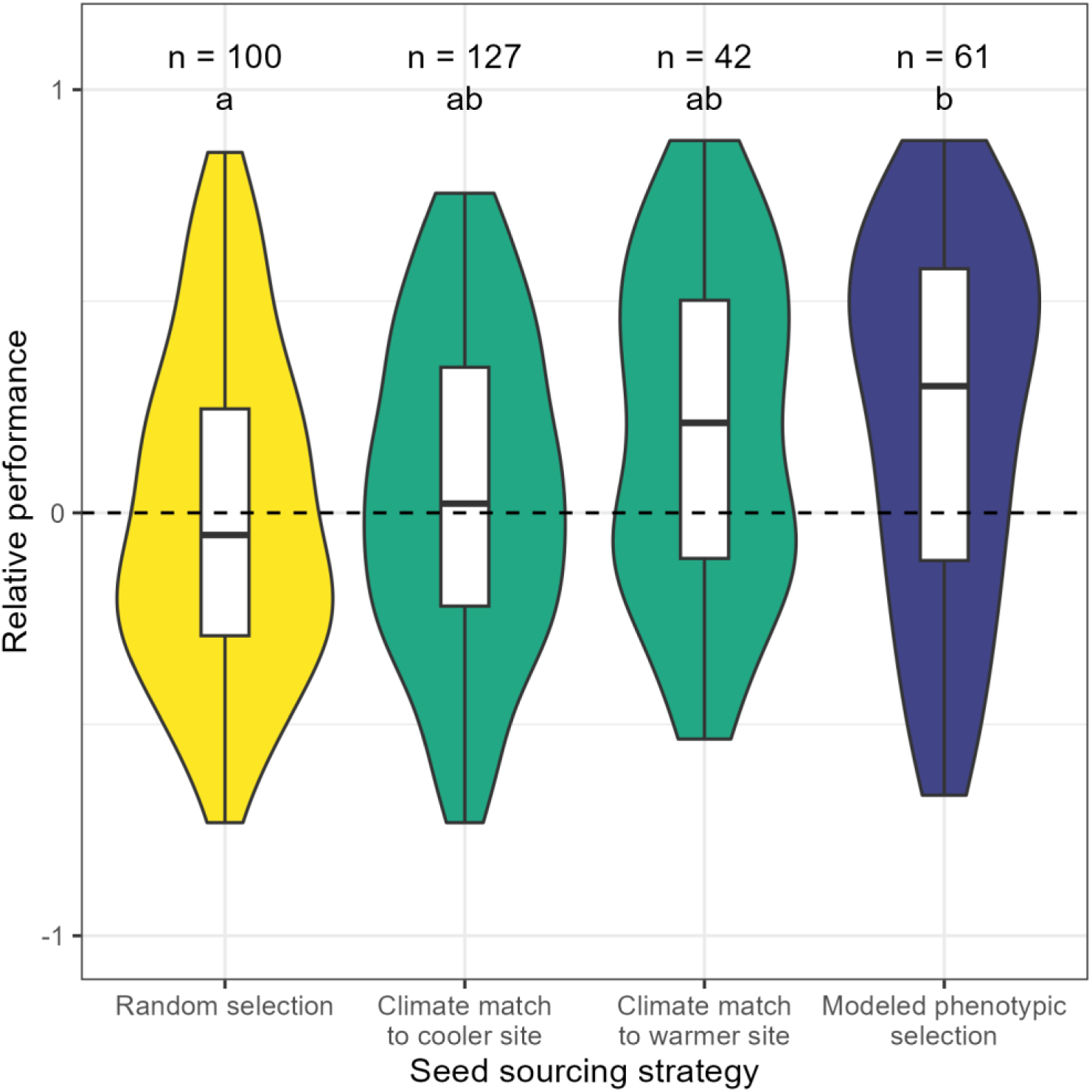
Violin plots (solid color) with overlayed boxplots of quartiles (white) showing comparison of seed sourcing outcomes modeled via simulated transfer of families from the cooler garden to the warmer garden. Performance metric is the family multiplicative fitness function measured in year 12 in the warmer garden, standardized to the garden-wide mean and range. A relative performance of 1 indicates the highest performance observed in the garden in year 12, whereas a relative performance of −1 indicates the lowest observed performance. The scale is centered around the garden-wide mean performance (dashed line). Random selection (yellow): 100 families selected at random. Climate match (green): families selected if origin site T_max_ is within 0.5°C of the 2014-2024 garden T_max_ (left: cooler, right: warmer). Modeled phenotypic selection (dark blue): selected families with 90^th^ percentile predicted growth at the cooler site based on the GAM fitted using year-5 data. Modeled phenotypic selection is the only strategy with significantly better performance than random selection.

## Discussion

Valley oak trees showed continued evidence of adaptation to climates of 21,000 years ago rather than present-day climates after ten years in two common gardens as first discovered for 5-year-old trees (Browne et al. 2019). Moreover, trees aged 5-12 demonstrates that maladaptation worsens as trees age and is exacerbated by weather extremes. These findings raise concern about the future health of oak forests in California, but our further analyses indicate that strategic management of oak populations may enhance their resilience and persistence. By modeling the relationship between temperature transfer distance and fitness of trees in the common garden, we can select the most successful potential seed sources for reforestation. Thus, despite dire predictions for the already maladapted valley oak’s susceptibility to projected future temperature increases, there is great potential to conserve and restore their populations *in situ*.

### Maladaptation to climate persists after twelve years of growth

While trees showed a pattern of local adaptation to winter temperatures, with peak growth rates observed in trees from similar origin climates to the gardens, the relationships between summer maximum temperatures and fitness proxies were instead highly directional and of higher magnitude than the relationships with winter temperature. Cumulatively and across years, trees from hotter, dryer, and more precipitation-seasonal climates tended to perform better than trees from cooler, wetter, and more temperature-seasonal climates. Cumulative performance and growth rate both showed negative relationships with annual maximum temperature transfer distance, with extrapolated trait optima outside of the current species range. Genetic variation between maternal tree lineages also accounted for variance in performance and significantly predicted between- and among-year variation in growth rates, demonstrating an important contribution of family-scale genetic variation to growth outcomes. Previous work on young seedlings in the same study also found that genomic estimated breeding values were more predictive of seedling success than source climate (Browne et al. 2019). Similarly, phenotypic differentiation among families, even those from similar climates, is consistent with other provenance studies (Savolainen et al. 2004, Wilczek et al. 2014, Lind et al. 2024). The decoupling of adaptation to temperature minima from that of temperature maxima may indicate that cold stress is a more influential selective pressure than heat or drought stress to the plant (thus allowing less-adapted phenotypes to persist amid post-glacial warming; Brady et al. 2019a). Regardless, evidence of maladaptation to summer temperature suggests meaningful and urgent vulnerability of the species even if local adaptation to other climatic conditions persists. Furthermore, we note that hot-summer origin sites tended to have moderate winter temperatures, possibly indicating that the apparent local adaptation was an artifact of an underlying maladaptive trend. We thus find strong, lasting evidence in a 12-year common garden study of current maladaptation to climate in valley oak, predicted by genetic and ecological characteristics of the maternal trees and their climates of origin.

Our results present evidence that valley oak trees are no longer present in the environments where they should theoretically grow best. Extrapolation of the temperature/growth rate relationships predicted that annual growth rates should peak at a temperature transfer distance of roughly −5°C, which is cooler than any observed transfer distance from a source site within the current valley oak species range. Likewise, given that the warmer garden does not represent a climatic extreme of the valley oak range (Delfino Mix et al. 2015), the hottest-on-average areas of the species range could heavily constrain growth of even the most warm-adapted valley oak genotypes if average and acute temperatures rise there. However, models predict that cooler inland parts of the species range will experience the most warming relative to historical conditions (Wang et al. 2016, AdaptWest Project 2022), potentially mediating vulnerability of warm-adapted genotypes and increasing vulnerability of cool-adapted genotypes. While we were unable to directly test the effects of precipitation on growth, we do note an anecdotal indication that changes in the temperature/growth relationship may be associated with annual variation in precipitation. As water availability has been shown to mediate valley oak response to temperature (Mead et al. 2019), it is likely that both temperature and precipitation will play a large role in determining valley oak fitness under future climates. Nonetheless, our findings demonstrate a clear and immediate danger of temperature increases not only to the least heat-adapted valley oak populations but to the entire species. As acute maximum temperatures increase over time, the range-wide negative effect of hot years will likely constrain growth of valley oaks in the aggregate (e.g., Sork et al. 2010).

An additional factor that gives trees a benefit in common gardens is the possibility of a longer growing season. As reported previously by Wright et al. (2021), progeny of trees from hotter environments had significantly earlier leaf emergence than trees from cooler environments. We found that earlier leaf emergence was correlated with higher growth rate and relative performance likely due to a longer growing season. Leaves also emerged earlier on average in years with warmer temperatures during the bud period (February-May), consistent with a previous study on valley oak (Gerst et al. 2017). In turn, we did not observe a tradeoff between a lengthened growing season and risk of frost damage; though late frosts occurred at the cooler garden, we did not observe negative fitness effects of early leaf emergence. This finding suggests a possible future adaptational lag; that is, trees adapted to environments cooler than future conditions may continue to wait longer than necessary to put out leaves and in turn reduce their growth potential. However, the effects of frost damage on other aspects of plant health (e.g., leaf morphology, biomass and nutrient allocation) should be considered as well. As we detected a pattern of local adaptation to mean winter temperature, other mechanisms likely impose a fitness penalty on families adapted to warmer environments than the garden. We thus find evidence that timing of leaf emergence partially predicts the fitness differences among valley oak trees from differing climates and may be a driver of future shifts toward lower fitness in cool-adapted populations.

### Maladaptation persists, and even worsens, as trees age

After a decade of growth in the common gardens, trees have shown limited to no acclimation to their planting locations; hot-source individuals continue to show significantly higher fitness than cool-source individuals, and the difference is more pronounced in many of the recent study years than early years. In addition, fitness continues to vary greatly with annual temperature maximum. The limited but apparent “leveling off” of fitness differences in 2022 and 2023 coincides with both years having maximum temperatures close to the 30-year average; the more pronounced relationship in 2024 confirms that hotter yearly temperature maxima still promote maladaptive patterns in valley oak. While phenotypic plasticity has been observed between trees in the two gardens (MacDonald 2017, Browne et al. 2019, Wright et al. 2021), and now additionally among years, individual plasticity has not been sufficient to overcome the underlying genotypes of the maternal trees. The decade-long persistence of climatic fitness differentiation demonstrates a clear increased risk among individuals across the species range, as genotypes are not locally adapted to their current environments and are thus even less adapted to predicted future climates. Any potential future acclimation would also depend on short-term climate, especially as average temperatures continue to rise. There is an inherent limitation to our ability to separate the effects of individual plant ontogeny and yearly climatic fluctuations in the gardens, but additional years of monitoring will allow for further decoupling of these factors especially as trees reach maturity. Likewise, our results only take temperature change into account; increased drought is also predicted to imperil valley oak populations (e.g., McLaughlin and Zavaleta 2012). Regardless, our findings after a decade of growth in the gardens suggest that early findings of maladaptation (Browne et al. 2019) were not restricted to the seedling stage and were not a result of maternal effects; maladaptive patterns have instead persisted, and even worsened, for over a decade.

### Plasticity in growth is insufficient as an “escape route” from maladaptation

We found strong signatures of interannual variation in metrics of valley oak fitness. In years with higher local temperatures at the gardens, negative relationships between summer temperature transfer distance and growth rates were more pronounced and growth rates were lower on average. The stronger negative relationship in hotter years supports the maladaptation hypothesis and suggests that populations in the hottest areas of the species range may be close to their climate threshold; in the hottest study year (2021), the temperature at the warmer garden was higher than all maternal seed source historical maxima (i.e., no individuals had temperature transfer distance < 0). In addition, the difference in maladaptive relationship between 2021 and all other years may indicate a threshold of warming tolerance. In most years, the predicted annual relative growth rates between 0°C and +5°C transfer distance were similar, but in 2021 the predicted growth rate was nearly 20% lower among individuals with transfer distance of +5°C. Therefore, we may see not only increased climate mismatch but also increased penalties for climate mismatch as the climate warms beyond the limits of trees’ ability to respond to changes. By contrast, the pattern of local adaptation to mean winter temperature seen in the cumulative analyses was also consistently detected within years. While there were some fluctuations in the relationship between winter temperature and growth (the peak around zero transfer distance was more pronounced in some years than others, and 2016 showed high growth in the cool-origin trees, similarly to the summer temperature relationship), the pattern stayed generally similar across the study period. However, while the best growth outcomes were observed in trees from origin sites with similar winter temperatures to the gardens, warmer-origin trees still consistently outperformed cooler-origin trees. Therefore, we detect an additional indication of underlying maladaptation. The negative relationships between winter temperature and growth are also slightly more pronounced in hotter study years, which may suggest that while valley oaks kept pace with changes in temperature since the Last Glacial Maximum, their adaptation may begin to lag under current and future conditions. The interannual fluctuations in growth rate also demonstrate individual phenotypic plasticity across years, but the nonsignificant effect of the family*time interaction indicates that the degree of plasticity is not genetically differentiated and thus not associated with climate of origin (Chambel et al. 2005). Furthermore, individual plasticity did not compensate for the genetic differentiation among families, as those from cooler origins consistently had lower growth despite interannual differences. Directional genetic differentiation overshadowing individual plasticity is consistent with the hypothesis that valley oak trees are lag-adapted as it points to historical local adaptation to varied climates as opposed to species-wide generalization through plasticity (Sultan 1995). We thus find that interannual fluctuations in temperature predict changes in valley oak growth rate over time.

Changes in the relationship between temperature transfer distance and fitness indicated a role of ontogeny in shaping individuals’ responses to climate fluctuations. In the third year of growth in the gardens the direction of the relationship between temperature transfer distance and growth rate reversed from other years. Initially, trees with cooler source climates than the gardens (i.e., positive transfer distance) had the fastest growth, followed by trees with similar source climates to the gardens (i.e., transfer distance around zero). After the fourth study year (second year after outplanting), trees with source climates cooler than the garden consistently grew less per year than those from hotter source climates and the signature of limited local adaptation was no longer present. The fitness difference between hotter-sourced trees and all other trees increased in magnitude for several years and has persisted for the remaining study years. The first two garden years were also the coolest years of the study period, suggesting that short-term fluctuations in local climate have a great degree of influence over short-term growth. However, if these fluctuations were the main driving force behind interannual growth variation we would expect the negative relationship to similarly reverse in subsequent cool years, but in fact the cooler-source trees consistently had lower fitness from 2017 onwards, even in years with similar temperature to 2015 and 2016. These early successes of cool-adapted and locally adapted individuals thus indicate a meaningful ontogenetic distinction within the first few years of growth in which young seedlings show different resource needs than even slightly older saplings (McLaughlin and Zavaleta 2012). The small local maximum growth rate around zero in the second year does suggest some degree of local adaptation; this may be evidence of maternal effects from locally adapted seed sources persisting into the seedling stage and conferring some degree of fitness benefit (Roach and Wulff 1987, Donelson et al. 2018). Indeed, maternal effects were shown to strongly predict early-stage seedling performance in another *Quercus* species (Gimeno et al. 2008). However, any benefit we observed was apparently short-lived (lasting only two years) and has not conferred a detectable lasting advantage to progeny in the gardens.

### Growth data from a provenance trial can improve seed transfer outcomes

We found that selection of high-performing phenotypes in a common garden significantly improved fitness outcomes of simulated seed transfer relative to random and climate-based selection. While valley oak showed consistent patterns of maladaptation across a decade of growth in the gardens in which warmer-sourced trees consistently outperformed cooler-sourced trees, some individuals performed better than others within and across years. We therefore identified 61 genotypes with high performance across gardens and over time, indicating the feasibility of climate-informed seed transfer to mitigate current and ongoing maladaptation. By incorporating phenotypic and genetic data, our hypothetical seed sourcing strategy was more effective than climate-based alternatives. Furthermore, the phenotypic criteria allow for selection of high-performing genotypes across a wider part of the species range than climate match strategies, which are limited by few seed source locations having temperatures in the recent past (1951-1980) similar to current or projected future temperatures of candidate planting sites. Indeed, there were no potential seed sources with past temperature ≥ 2°C higher than the warmer garden, so no “future match” scenario was possible in our analysis. Given that valley oak is already growing suboptimally under current climates, it is unsurprising that a climatic approach to seed sourcing could quickly run up against the upper temperature limit of the species range. By contrast, the empirical common garden data confirms that fitness varies within populations and high-performing individuals are present even in the more temperate parts of the species range, and some variation in success is due to genetic variation as opposed to local adaptation. This is consistent with other studies that report a high importance of existing genetic variation for the success of assisted gene flow (e.g. Beaulieu and Rainville 2005, Robert et al. 2024). Our results also broadly agree with those of Browne et al. (2019), who used the same common garden study to predict high-performing genotypes using genomic tools and determined that selection based on genetic markers can improve seed sourcing outcomes. This concordance indicates that empirical provenance data can effectively validate genomic methods, which has been a central goal of recent research on local (mal)adaptation in the context of global change (Huxman et al. 2022, Schwinning et al. 2022). In turn, our findings show that it is possible to retain the valley oak ecosystem in place as opposed to taking more drastic approaches like replacing them with other more climate-resilient oak species.

### Five-year seedling studies provide sufficient data to predict twelve-year maladaptive patterns

Predictions based on data collected in the fifth year of the study (the third year after outplanting in the gardens) were sufficient to (a) robustly identify maladaptive patterns and (b) find high-performing genotypes for transfer into warmer conditions. While the earliest study years did show single-year variation in outcomes (most notably, cool-origin trees grew more quickly than warm-origin trees in the first two years following outplanting), the overall relationship between source climate and growth observed in year 5 did not change directionality for the remainder of the study to date. Early differences in observed adaptation may have been caused by either maternal effects on seedlings (Roach and Wulff 1987, Gimeno et al. 2008, Donelson et al. 2018), outplanting effects (Close et al. 2005), or anomalous low temperatures (2015 and 2016 were the two coolest years in the study period). These early anomalies were only detectable when examining single growing seasons; by contrast, cumulative metrics were consistently indicative of a negative relationship between temperature and growth. Therefore, we suggest that seedling studies can be effective tools for climate-adaptive restoration if longer-term studies are not feasible. The question remains whether the fitness differences among cohorts may eventually decrease, given the long lifespan of valley oak trees. We acknowledge that our measured fitness outcomes are in 12-year-old trees so our findings may not reflect patterns in adult life stages, especially in natural as opposed to controlled garden conditions. Fitness differences in provenance studies can be dependent on temporal scale--for instance, Germino et al. (2019) only observed local adaptation after 14 years of growth--but other studies (e.g. Kashian and Barnes 2021, Chmura and Modrzyński 2023) found short-term patterns in adaptation to remain consistent across decades. Regardless, early growth years are important for successful tree recruitment and establishment (e.g., Snyder and Ellner 2022), and valley oak has been shown to suffer from limited recruitment and survival into adulthood (Tyler et al. 2006), so consistent maladaptation in years 0-12 is likely to impose a lasting fitness cost even if performance changes in later growth stages. In turn, these early data can be applied to improve seed selection for reforestation. The efficacy of relatively short-duration studies combined with the broad agreement between genomic and common garden approaches presents powerful evidence that better-informed revegetation projects are within reach even with limited resources. Future studies can corroborate common garden results to genomic markers in wild populations to promote more informed predictions of (mal)adaptive dynamics in wild populations (see Lind and Lotterhos 2024). Crucially, we show that phenotypically based seed selection is “worth the effort” relative to selection based solely on climate projections.

## Conclusions

Valley oak as a species shows tremendous vulnerability to higher temperatures in the future, given that the species is maladapted to current temperatures. Interannual fitness variation among genetic lineages illustrates the sensitivity of tree response to annual climate variation and confirms range-wide maladaptation. Nonetheless, high-performing individuals with warm-adapted phenotypes are still present within populations across the species range, especially but not only in warm environments. This pattern of genetically determined high-performers despite prevailing maladaptation indicates feasibility of restoration via genotype-informed seed transfer. Our findings foretell that in the absence of active intervention, maladaptation will first negatively impact individuals in the cooler part of the species range, but the entire species will show reduced growth due to increasing temperatures. Furthermore, the relative fitness penalty of maladaptation may increase under warmer climates. Therefore, revegetation of the species using seeds preadapted to future conditions provides an encouraging path forward for conservation of the valley oak savanna ecosystem.

## Author contributions

VLS and JWW designed the common gardens; VLS, JWW, and ARBG designed research questions; all authors performed research; MO contributed data from dissertation; ARBG analyzed data with input from VLS; ARBG and VLS wrote the paper with input from JWW.

## Acknowledgements

We recognize the native peoples of California as the traditional stewards of the ecosystems in which this research was conducted. We thank Annette Delfino-Mix, Courtney Canning, and Lisa Crane for their assistance in establishing and maintaining the provenance trial. We particularly thank Paul Gugger who led the effort and organized the teams to collect the 11,000 acorns to start the provenance study, as well as numerous past and present Sork Lab members and volunteers who assisted with data collection. This research was funded by an NSF long-term research grant awarded to VLS and JWW (LTREB-2232794), by funds from the USDA Forest Service Pacific Southwest Research Station, and support to VLS from UCLA.

The findings and conclusions in this publication are those of the authors and should not be construed to represent any official USDA or U.S. Government determination or policy.

Any use of product names is for informational purposes only and does not imply endorsement by the US Government.

## Notes

### Competing Interest Statement

The authors have declared no competing interest.

