## Supplemental Material for "Current maladaptation increases with age in valley oak (*Quercus lobata*): Implications for future populations"

Supplementary information text

Materials and methods

Principal components analysis on maternal origin climate data

Data on maternal tree source site climates from the California Basin Characterization Model (Flint et al. 2013) were used to compare common garden growth outcomes against historical climatic conditions experienced by the maternal source trees. All measurements were 30-year averages of 1950-1981 data gridded at 270m pixel resolution. Temperature and precipitation data were used to calculate the BioClim variables (worldclim.org), and a principal components analysis was run on 10 climate variables (Maximum summer temperature, minimum winter temperature, maximum, minimum, and average annual temperatures, climatic water deficit, temperature seasonality, precipitation seasonality, precipitation of warmest quarter, precipitation of coldest quarter). The first PC axis (40.8% of variance explained) represented the spectrum between hotter, dryer sites and cooler, wetter sites, while the second PC axis (23.3% of variance explained) represented seasonal differences in temperature and precipitation (higher values indicate temperature-based seasonality, lower values indicate precipitation-based seasonality). For more detail on the climatic variables and PCAs, see Browne et al. 2019.

Testing for maladaptation: longitudinal cohort study

To explicitly test for climatic maladaptation (e.g., families from warmer source sites than the garden showing higher fitness than families from cooler or similar-temperature source sites), we binned families into five transfer distance cohorts and compared their MFF over time. Transfer distance values were calculated across all observations using the median summer maximum temperature within the study period (cooler garden: 2019, warmer garden: 2023; see Table S12). Cohorts were defined to represent discrete bins of transfer distance values, generally but not always in 2° C increments, to capture local modal transfer distance values as much as possible while also considering relative sample sizes (Figure S2). We then conducted repeated-measures ANOVAs of the cohort*year interaction on MFF with family as a random factor using the aov() function in R package stats. Gardens were analyzed separately since the cohorts were garden-specific. To validate the cohort groupings, we tested several series of similar cohort definitions, including random subsets of individuals. We consistently observed similar outputs, suggesting that the exact cohort definitions do not substantially influence the results. Note that these are not the same cohorts used to illustrate growth rate differences among years.

Effects of geography on growth and fitness

We tested the effects of geography on growth rate and the multiplicative fitness function (MFF). As we were interested in detecting geographic “hotspots” of overall successful families, we measured response of cumulative MFF and growth rate (MFF in 2023; 2014-2024 cumulative growth rate) to latitude, longitude, and the first two principal components describing variation in maternal seed source sites (higher PC1: hotter/dryer, higher PC2: temperature, not precipitation, drives seasonality; see “Maternal tree source climate data” section above). Gardens were analyzed separately. The growth rate models also included the fixed effects of initial height and garden block and the random effect of family. Explanatory variables were scaled as Z-scores. Phenology and geography models were fitted using the lmer() function in R package lme4 (Bates 2010).

Finally, to visualize and interpret the geographic differences in MFF, we used two-dimensional kriging to interpolate MFF values between points within the hypothesized species range. Models were fitted using the gstat() function, and interpolated using the interpolate() function, in R package terra version        1.7-83 (Hijmans et al. 2024).

Results and discussion

Testing for maladaptation: longitudinal cohort study

To answer Q1 and Q2, we used a cohort approach to test whether individuals with similar source temperatures provided evidence of maladaptation. If populations are maladapted due to lag-adaptation to cooler climates, we expect that families from hotter climates would outperform those from cooler climates. We found significant effects of cohort, year, and cohort*year on MFF of individuals across both gardens (cooler garden: *F*_cohort, df = 4_ = 11.2, *p <* 0.001; *F*_year, df=8_ = 55.7, *p <* 0.001; *F*_cohort*year, df=32_ = 9.5, *p <* 0.001; warmer garden: *F*_cohort, df = 4_ = 2.9, *p* = 0.02; *F*_year, df = 6_ = 76.2, *p <* 0.001; *F*_cohort*year, df = 24_ = 3.6, *p <* 0.001). Tracking MFF of cohorts of individuals (grouped based on their temperature transfer distance in the year with median garden temperature) across the study period also demonstrated 2015-2016 as an inflection point for the emergence of maladaptation; the pattern persisted for the remainder of the study period and the fitness difference between the trees from the hottest and coolest source populations increased over time (Figure S7). In the cooler garden, cohorts B (source site 0-2 degrees C hotter than garden) had the highest fitness in 2015 and a large drop in 2016, after which cohort A (source site 2+ degrees hotter than garden) had consistently higher fitness. Cohorts A, C, and D had relatively similar fitness from 2014-2017, but cohort E consistently had the lowest fitness even in the initial years. Fitness differences among cohorts were more variable across years in the warmer garden than in the cooler garden. Generally, cohorts A and B (cumulatively, source site between 2 degrees hotter and 2 degrees cooler than garden) had the highest fitness across most years and the remaining 3 cohorts had similarly low fitness across years. As in the cooler garden, cohort E (source site 7+ degrees cooler than garden) consistently showed the lowest fitness. In contrast to the cooler garden, the warmer garden showed more consistent fitness responses of all cohorts across years (e.g., all cohorts showed an increase from 2017-2018 and a decrease from 2020-2021). The random effect of maternal lineage explained much more of the variance than the fixed effects (cooler garden: marginal R^2^ = 0.09, conditional R^2^ = 0.75; warmer garden: marginal R^2^ = 0.06; conditional R^2^ = 0.72). Full model outputs are provided in Tables S13 and S14. The two gardens provide two tests of the maladaptation (Q1), but it is important to note that cohorts were independently defined for each garden and do not represent the same maternal tree individuals across both gardens.

Effects of geography on growth and fitness

To test for the effect of geographic variation, including photoperiod, on cumulative MFF and growth rate, we tested the relationship between 2024 MFF/2014-2024 growth rate and longitude/latitude when controlling for the maternal climate PC vectors. We did not find significant relationships between MFF and latitude or longitude (no macroclimate effect), but the PC vectors, representing local climatic variation, significantly predicted MFF. Families from hotter, dryer, and more temperature-seasonal (as opposed to precipitation-seasonal) sites had significantly higher fitness (cooler garden: *p*_PC1_ < 0.001, *p*_PC2_ = 0.01; warmer garden: *p*_PC1_ = 0.02, *p*_PC2_ < 0.001). Geographic variation was high even at the local climate scale, with trees from nearby sites showing varied fitness outcomes. Several maternal trees in the 95^th^ percentile for MFF were sourced from cooler sites than the gardens and many sites hotter than the gardens did not have high performing trees (Figure S8). Geographic effects on cumulative growth rates were slightly different. At the cooler garden, latitude and PC1 both significantly predicted growth rate (*p*_latitude_ = 0.03, *p*_PC1_ < 0.001) as higher in families from hotter sites in the northern extent of the species range. At the warmer garden, only PC2 significantly predicted growth rate (*p*_PC2_ = 0.03), and families from more precipitation-seasonal sites had higher growth rate. Full model summaries are provided in Tables S15-S18.

We did not find strong patterns of latitude or longitude predicting fitness such as reported for forest trees in Canada (e.g., Rehfeldt et al. 1999), nor did individuals clearly separate into distinct geographic units (Liao et al. 2022). Instead, we found high geographic variation in maternal tree fitness. Furthermore, though local climate was a significant predictor of MFF, several trees in the 95^th^ percentile for MFF were from sites up to 2 degrees cooler than the median garden temperature.

Supplementary figures

Figure S1: Total annual precipitation 2014-2024 in the warmer garden (red) and cooler garden (blue), with the mean of both gardens shown in the green dashed line. Data from PRISM (prism.oregonstate.edu).

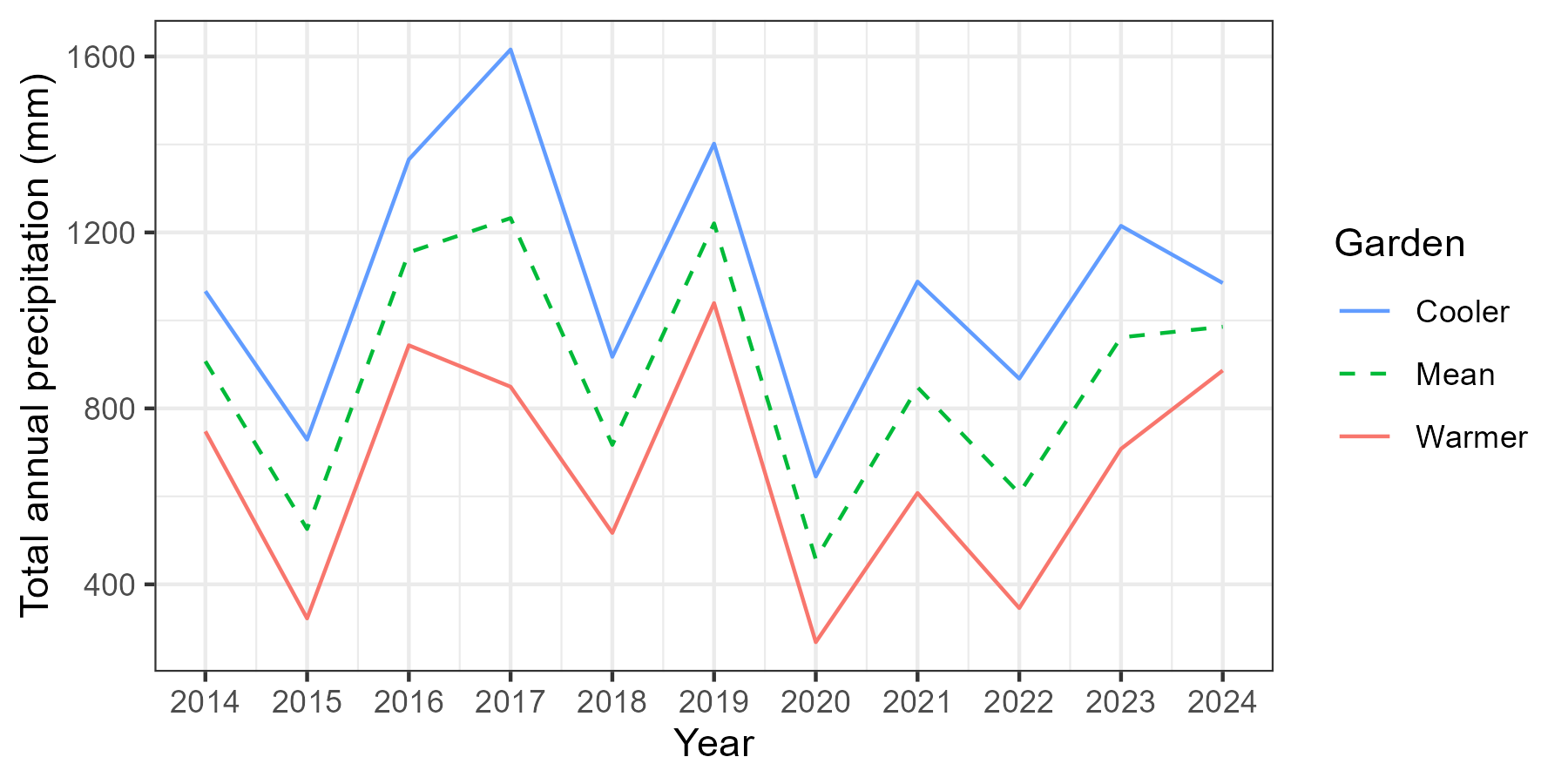

Figure S2: Ten-year cumulative trends in growth rate (left) and performance (right) as a function of temperature transfer distance (maximum monthly temperature averaged across June-August) between maternal location and common garden. Solid curves denote generalized additive models fit to empirical data, and dashed curves denote extrapolation of the models using Gaussian process regression. Shaded regions represent 95% confidence intervals. The peak in each extrapolated curve represents the theoretical species-wide trait optimum. Vertical dashed line in each plot denotes where source temperature equals garden temperature; vertical solid lines denote estimated average temperature during the Last Glacial Maximum (21 kya; Wang et al. 2012) and predicted average temperature in year 2100 under a stabilization climate scenario (RCP 4.5; AdaptWest Project 2022). The area to the left of the zero line contains individuals from warmer historical climates than the garden, while the area to the right contains individuals from cooler historical climates.

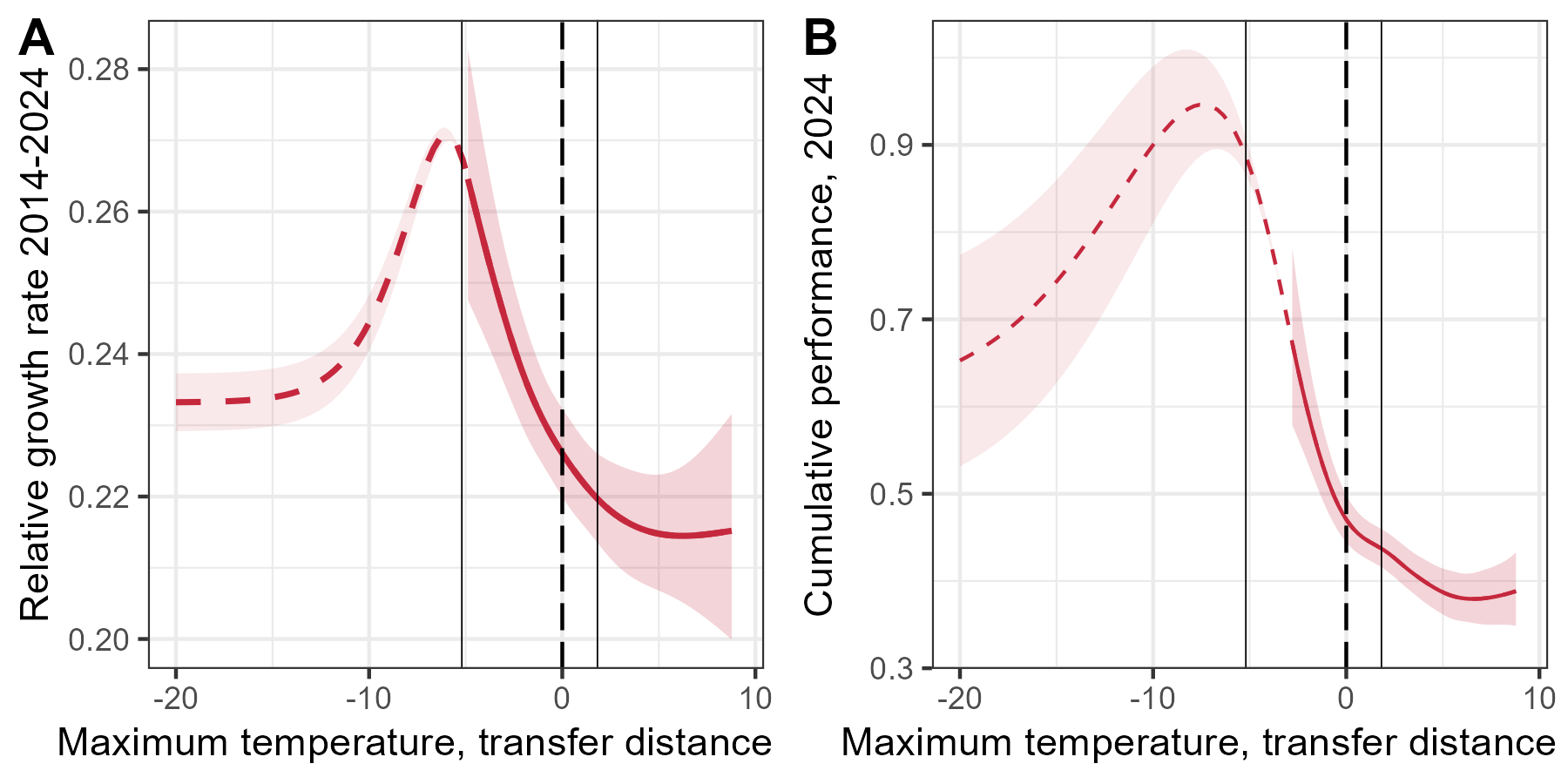

Figure S3: Yearly trends in valley oak growth rate as a function of transfer distance between maternal location and common garden, arranged and color-coded in order of cooler to warmer temperatures. Solid colored curves represent empirically derived generalized additive model predictions of single-year relative growth rate as a function of yearly temperature transfer distance (maximum monthly temperature averaged across June-August) across the period of tree growth in the common gardens, 2015-2024 (2018 is excluded due to incomplete data). Long-dashed curves represent extrapolations of the model using Gaussian process regression. Thin dotted curves represent non-significant relationships; thicker dotted curves represent marginally significant relationships (p < 0.1). Vertical dashed line in each plot denotes where source temperature equals garden temperature; vertical solid lines denote estimated average temperature during the Last Glacial Maximum (21 kya; Wang et al. 2012, Karger et al. 2017) and predicted average temperature for years 2070-2100 under a stabilization climate scenario ensemble (RCP 4.5; AdaptWest Project 2022). The area to the left of the zero line contains individuals from warmer historical climates than the garden, while the area to the right contains individuals from cooler historical climates.

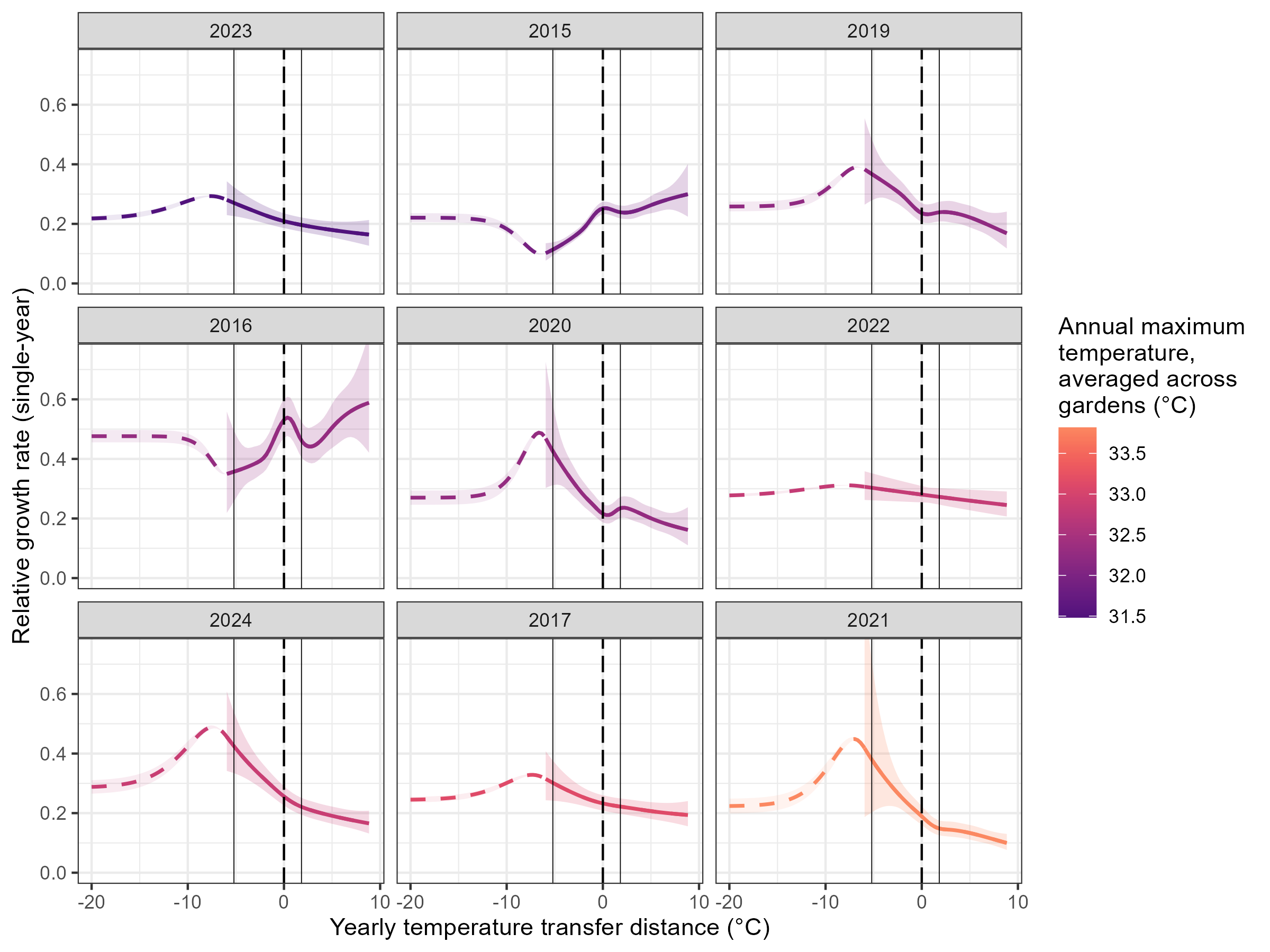

Figure S4: Distribution of maternal tree origin site temperatures (summer maximum; winter mean, °C), colored by elevation above sea level (m). Pearson correlation coefficient between summer maximum and winter mean temperature = -0.01. An elevational cline effect is apparent among sites with low summer and winter temperature (bottom right corner); low-elevation sites show wide variation in temperature. Note that all origin sites with summer maximum temperature above 33°C have winter mean temperature between 7 and 10°C. Temperature data reflects 1951-1980 averages sourced from the PRISM database.

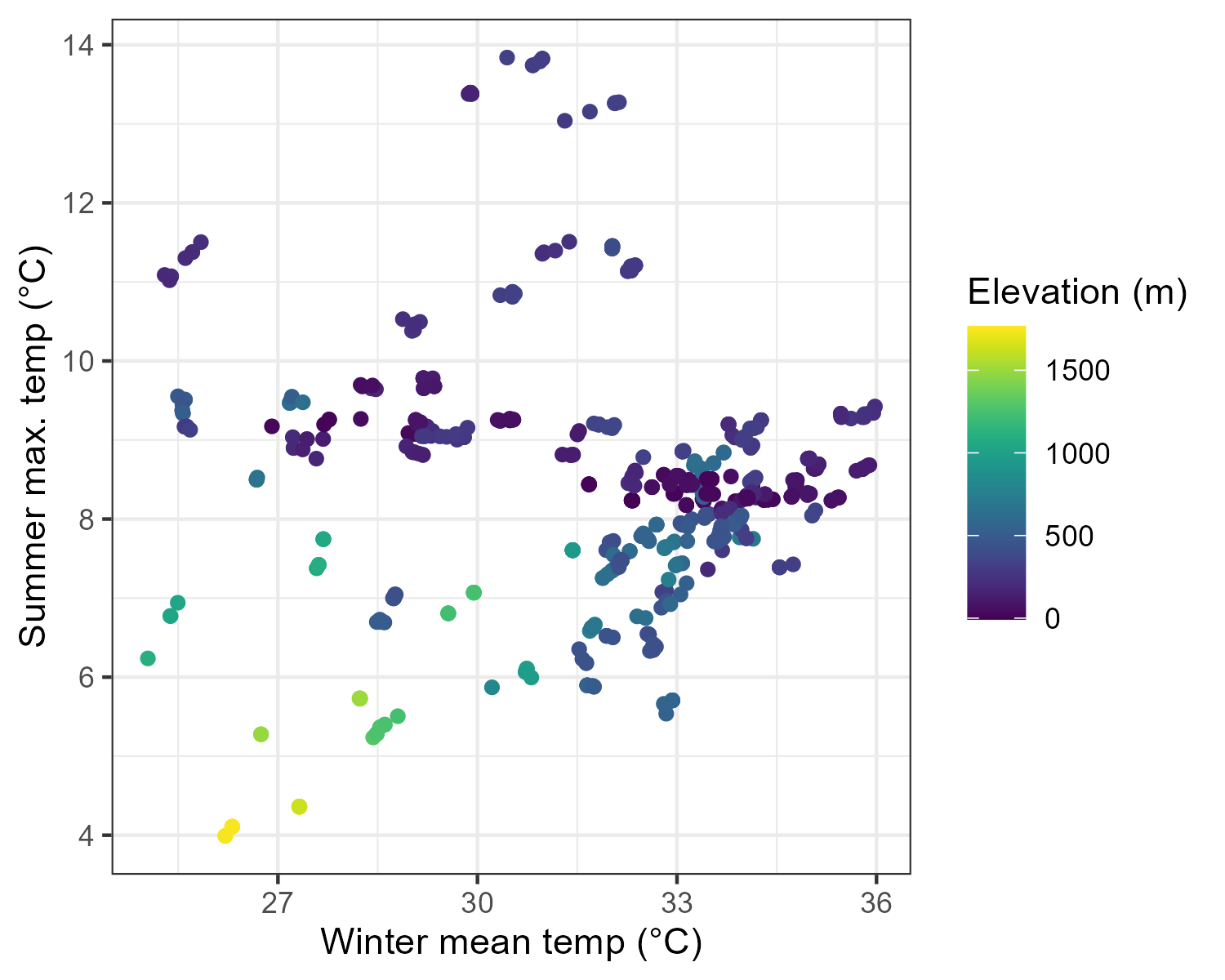

Figure S5: Comparison of quantile cutoffs for modeled phenotypic selection (selected based on performance in year 5 at the cooler garden, outcome based on performance in year 12 at the warmer garden). We used the 90th percentile in our analysis, but patterns are similar across percentiles 75-90. 50th-percentile selection (left, shown for comparison) is much lower than the others but still shows a median value > 0 relative performance. As expected, we observe an incremental tradeoff between number of phenotypes and median performance as the cutoff increases. However, selecting only the 95th percentile (right) of phenotypes in year 5 actually results in slightly lower median performance and a 50% reduction in sample size compared to the 90th percentile.

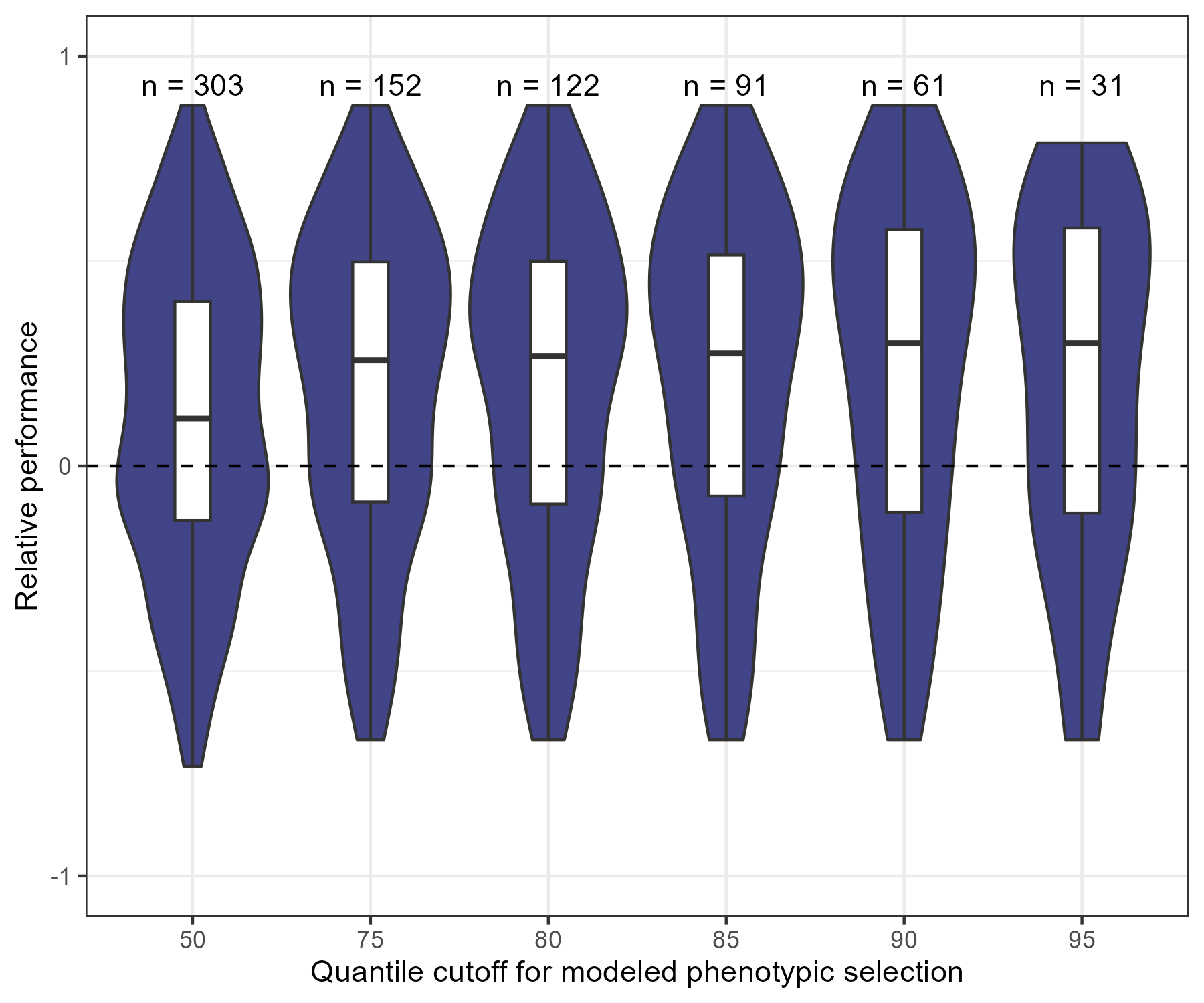

Figure S6: Distribution of (garden – site) temperature transfer distance values from which cohorts were derived for analysis for the cooler garden (top) and warmer garden (bottom). Bars represent counts of individuals, colors indicate cohorts.

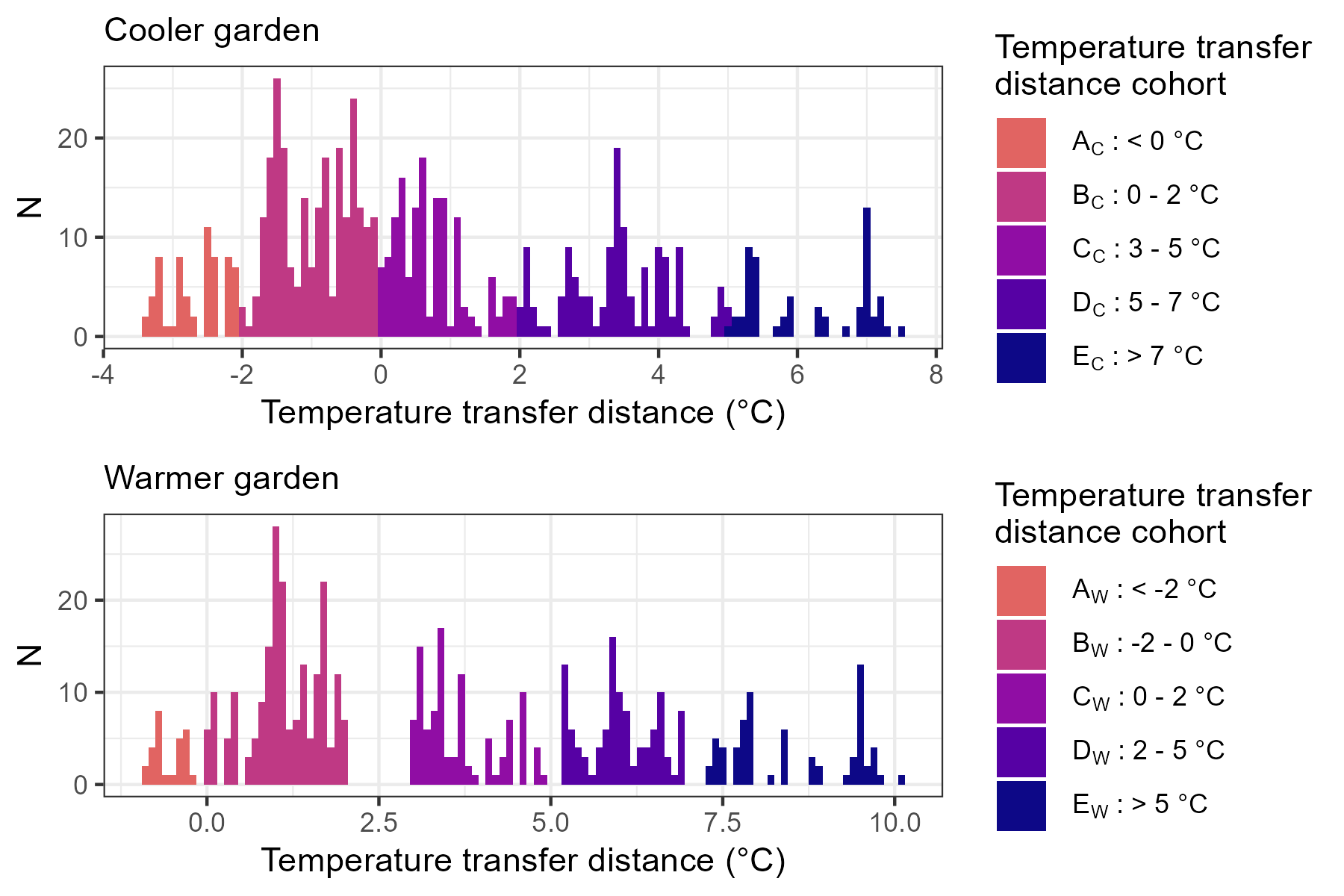

Figure S7: Means and standard errors of performance of five tree family cohorts at each garden. Differences in MFF by year, cohort, and the year*cohort interaction were all statistically significant (Tables S7 and S8). Each cohort was assigned based on yearly temperature transfer distance between source and garden (e.g., cohort AW consists of plants growing in gardens at least 2°C cooler than their source locations) in the year with median garden temperature (cooler garden: 2019, warmer garden: 2023).

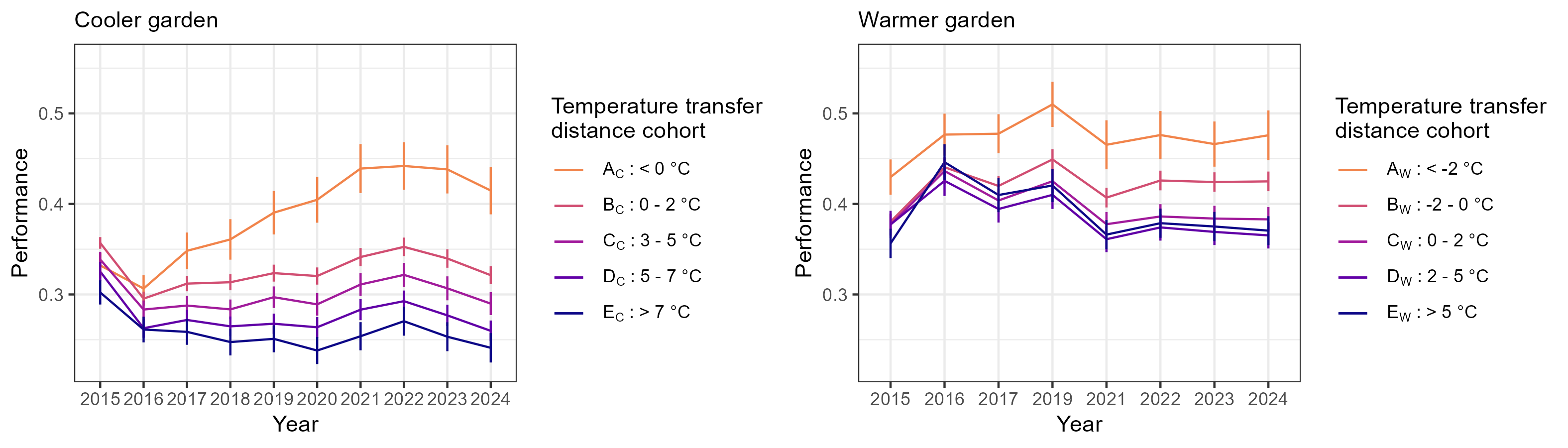

Figure S8: Geographic distribution of performance of maternal seed sources measured in the common gardens (left: cooler garden, right: warmer garden) with point locations indicated by blue or red dot surrounded by 2500-meter interpolated buffer zones. Color gradient of buffer zone shows family mean MFF in the common gardens in 2023. The figures also include dots for families from locations with temperatures that are cooler (blue) or hotter (red) than median of maximum summer temperatures in each garden. Black triangles show locations of maternal trees with 95th percentile MFF in each garden as of 2023.

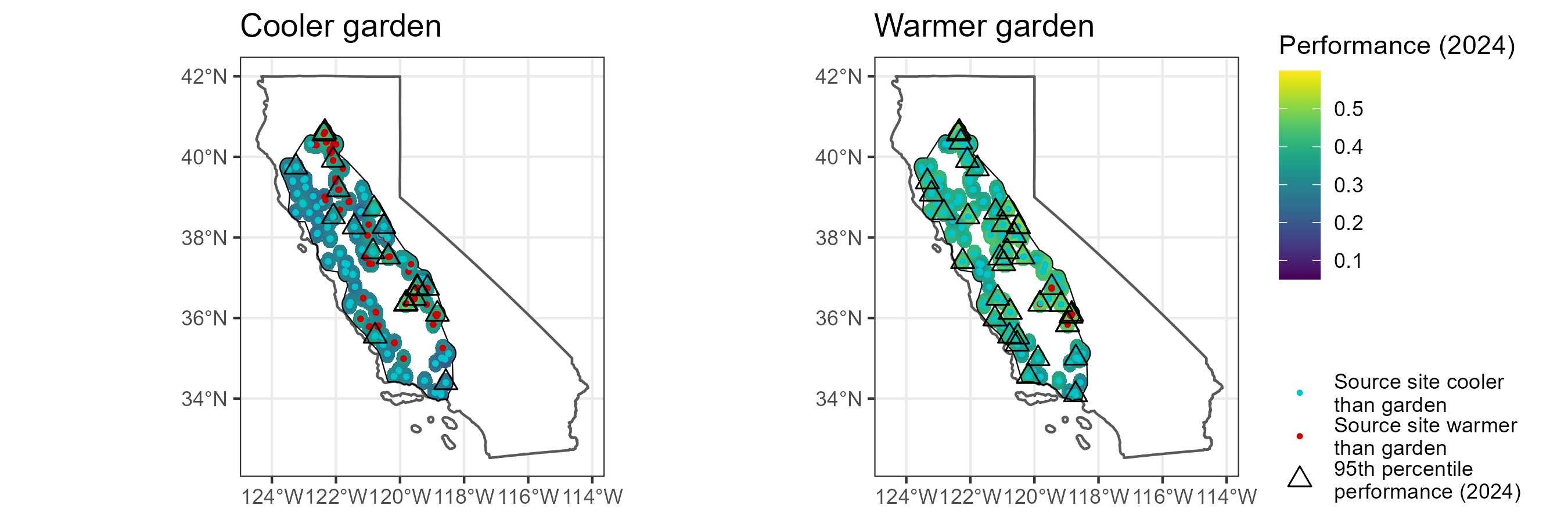

Supplementary tables

Table S1: Summary of growth variables measured on valley oak progeny in the common gardens in each year. Starting in 2018, some individuals were too tall to efficiently measure to their full heights so cutoff thresholds (shown in parentheses) were imposed.

| **Year** | **Growth variables measured** |
| --- | --- |
| 2013 | Height, basal diameter |
| 2014 | Height, basal diameter |
| 2015 | Height, basal diameter |
| 2016 | Height |
| 2017 | Height |
| 2018 | Height (<4 m), DBH (haphazard subset), basal diameter (haphazard subset) |
| 2019 | Height (<4 m), DBH (all trees above 150 cm height) |
| 2020 | DBH or basal diameter (IFG: all; CSO: 4/5 blocks) |
| 2021 | Height (<3 m), DBH (all trees above 150 cm height); basal diameter (removed trees), actual height (removed trees) |
| 2022 | Height (<2 m), DBH (all trees above 150 cm height) |

Table S2: Full model summary for GAM estimating cumulative effect of mean winter temperature transfer distance on relative growth rate, 2014-2024. Planting block, nested within garden, was included to account for unintended environmental differences within gardens but is not shown here for easier interpretation.

| **Component** | **Term** | **Estimate** | **Std Error** | **t-value** | **p-value** |  |
| --- | --- | --- | --- | --- | --- | --- |
| A. parametric coefficients | (Intercept) | -1.46 | 0.01 | -107.22 | 0.00 | *** |
|  | SiteBlockChico2 | -0.07 | 0.02 | -3.29 | 0.00 | ** |
|  | SiteBlockChico3 | -0.13 | 0.02 | -7.24 | 0.00 | *** |
|  | SiteBlockChico4 | -0.16 | 0.02 | -9.00 | 0.00 | *** |
|  | SiteBlockChico5 | -0.11 | 0.02 | -6.06 | 0.00 | *** |
|  | SiteBlockIFG1 | -0.22 | 0.02 | -10.88 | 0.00 | *** |
|  | SiteBlockIFG2 | -0.25 | 0.02 | -12.73 | 0.00 | *** |
|  | SiteBlockIFG3 | -0.27 | 0.02 | -13.64 | 0.00 | *** |
|  | SiteBlockIFG4 | -0.26 | 0.02 | -13.09 | 0.00 | *** |
|  | SiteBlockIFG5 | -0.35 | 0.02 | -16.52 | 0.00 | *** |
| **Component** | **Term** |  |  |  |  |  |
| B. smooth terms | Smooth term (2014 height) | 5.05 | 6.30 | 370.25 | 0.00 | *** |
|  | te(tdiff_win) | 3.02 | 3.41 | 7.15 | 0.00 | *** |
|  | Smooth term (Locality) | 55.21 | 94.00 | 1.56 | 0.00 | *** |
|  | Smooth term (Family) | 0.00 | 625.00 | 0.00 | 0.89 |  |
| Signif. codes: 0 <= '***' < 0.001 < '**' < 0.01 < '*' < 0.05 | | | | | | |
| Adjusted R-squared: 0.599, Deviance explained 0.545 | | | | | | |
| fREML : -2583.130, Scale est: 0.0133, N: 3351 | | | | | | |

Table S3: Full model summary for GAM estimating cumulative effect of mean winter temperature transfer distance on multiplicative fitness function, 2014-2024.

| **Component** | **Term** | **Estimate** | **Std Error** | **t-value** | **p-value** |  |
| --- | --- | --- | --- | --- | --- | --- |
| A. parametric coefficients | (Intercept) | -0.87 | 0.02 | -41.57 | 0.00 | *** |
|  | Garden: IFG | -0.30 | 0.03 | -10.62 | 0.00 | *** |
| **Component** | **Term** |  |  |  |  |  |
| B. smooth terms | s(tdiff_win) | 7.04 | 7.80 | 3.82 | 0.00 | ** |
|  | Smooth term (2014 performance) | 2.44 | 3.08 | 9.98 | 0.00 | *** |
|  | Smooth term (Family) | 0.00 | 621.00 | 0.00 | 0.56 |  |
|  | Smooth term (Locality) | 55.56 | 94.00 | 1.59 | 0.00 | *** |
| Signif. codes: 0 <= '***' < 0.001 < '**' < 0.01 < '*' < 0.05 | | | | | | |
| Adjusted R-squared: 0.236, Deviance explained 0.276 | | | | | | |
| fREML : -47.543, Scale est: 0.0476, N: 1229 | | | | | | |

Table S4: Full model summary for GAM estimating cumulative effect of annual maximum temperature transfer distance on relative growth rate, 2014-2024. Planting block, nested within garden, was included to account for unintended environmental differences within gardens but is not shown here for easier interpretation.

| **Component** | **Term** | **Estimate** | **Std Error** | **t-value** | **p-value** |  |
| --- | --- | --- | --- | --- | --- | --- |
| A. parametric coefficients | (Intercept) | -1.44 | 0.01 | -107.68 | 0.00 | *** |
|  | SiteBlockChico2 | -0.07 | 0.02 | -3.31 | 0.00 | *** |
|  | SiteBlockChico3 | -0.13 | 0.02 | -7.35 | 0.00 | *** |
|  | SiteBlockChico4 | -0.16 | 0.02 | -9.00 | 0.00 | *** |
|  | SiteBlockChico5 | -0.11 | 0.02 | -6.11 | 0.00 | *** |
|  | SiteBlockIFG1 | -0.26 | 0.02 | -12.95 | 0.00 | *** |
|  | SiteBlockIFG2 | -0.30 | 0.02 | -14.85 | 0.00 | *** |
|  | SiteBlockIFG3 | -0.32 | 0.02 | -15.76 | 0.00 | *** |
|  | SiteBlockIFG4 | -0.30 | 0.02 | -15.11 | 0.00 | *** |
|  | SiteBlockIFG5 | -0.39 | 0.02 | -18.42 | 0.00 | *** |
| **Component** | **Term** |  |  |  |  |  |
| B. smooth terms | Smooth term (2014 height) | 5.24 | 6.53 | 361.05 | 0.00 | *** |
|  | Smooth term (Tdiff, summer max) | 2.94 | 3.39 | 14.15 | 0.00 | *** |
|  | Smooth term (Locality) | 49.95 | 94.00 | 1.21 | 0.00 | *** |
|  | Smooth term (Family) | 0.00 | 625.00 | 0.00 | 0.88 |  |
| Signif. codes: 0 <= '***' < 0.001 < '**' < 0.01 < '*' < 0.05 | | | | | | |
| Adjusted R-squared: 0.600, Deviance explained 0.545 | | | | | | |
| fREML : -2595.198, Scale est: 0.0133, N: 3351 | | | | | | |

Table S5: Full model summary for GAM estimating cumulative effect of annual maximum temperature transfer distance on multiplicative fitness function, 2014-2024.

| **Component** | **Term** | **Estimate** | **Std Error** | **t-value** | **p-value** |  |
| --- | --- | --- | --- | --- | --- | --- |
| A. parametric coefficients | (Intercept) | -0.85 | 0.02 | -45.49 | 0.00 | *** |
|  | Garden: IFG | -0.35 | 0.03 | -13.45 | 0.00 | *** |
| **Component** | **Term** |  |  |  |  |  |
| B. smooth terms | Smooth term (Tdiff, summer max) | 3.60 | 3.88 | 15.56 | 0.00 | *** |
|  | Smooth term (2014 performance) | 2.45 | 3.10 | 9.14 | 0.00 | *** |
|  | Smooth term (Family) | 0.00 | 621.00 | 0.00 | 0.59 |  |
|  | Smooth term (Locality) | 44.53 | 94.00 | 0.97 | 0.00 | *** |
| Signif. codes: 0 <= '***' < 0.001 < '**' < 0.01 < '*' < 0.05 | | | | | | |
| Adjusted R-squared: 0.239, Deviance explained 0.268 | | | | | | |
| fREML : -67.180, Scale est: 0.0481, N: 1229 | | | | | | |

Table S6: Full model summary for GAM estimating cumulative effect of averaged summer maximum temperature transfer distance on relative growth rate, 2014-2024. Planting block, nested within garden, was included to account for unintended environmental differences within gardens but is not shown here for easier interpretation.

| **Component** | **Term** | **Estimate** | **Std Error** | **t-value** | **p-value** |  |
| --- | --- | --- | --- | --- | --- | --- |
| A. parametric coefficients | (Intercept) | -1.44 | 0.01 | -106.14 | 0.00 | *** |
|  | SiteBlockChico2 | -0.07 | 0.02 | -3.30 | 0.00 | *** |
|  | SiteBlockChico3 | -0.13 | 0.02 | -7.37 | 0.00 | *** |
|  | SiteBlockChico4 | -0.16 | 0.02 | -9.01 | 0.00 | *** |
|  | SiteBlockChico5 | -0.11 | 0.02 | -6.12 | 0.00 | *** |
|  | SiteBlockIFG1 | -0.27 | 0.02 | -13.05 | 0.00 | *** |
|  | SiteBlockIFG2 | -0.30 | 0.02 | -14.91 | 0.00 | *** |
|  | SiteBlockIFG3 | -0.32 | 0.02 | -15.80 | 0.00 | *** |
|  | SiteBlockIFG4 | -0.31 | 0.02 | -15.17 | 0.00 | *** |
|  | SiteBlockIFG5 | -0.40 | 0.02 | -18.43 | 0.00 | *** |
| **Component** | **Term** |  |  |  |  |  |
| B. smooth terms | Smooth term (2014 height) | 5.25 | 6.55 | 359.98 | 0.00 | *** |
|  | Smooth term (Tdiff, summer averaged max) | 2.91 | 3.38 | 14.00 | 0.00 | *** |
|  | Smooth term (Locality) | 50.28 | 94.00 | 1.22 | 0.00 | *** |
|  | Smooth term (Family) | 0.00 | 625.00 | 0.00 | 0.88 |  |
| Signif. codes: 0 <= '***' < 0.001 < '**' < 0.01 < '*' < 0.05 | | | | | | |
| Adjusted R-squared: 0.600, Deviance explained 0.545 | | | | | | |
| fREML : -2595.103, Scale est: 0.0133, N: 3351 | | | | | | |

Table S7: Full model summary for GAM estimating cumulative effect of averaged summer maximum temperature transfer distance on multiplicative fitness function, 2014-2024.

| **Component** | **Term** | **Estimate** | **Std Error** | **t-value** | **p-value** |  |
| --- | --- | --- | --- | --- | --- | --- |
| A. parametric coefficients | (Intercept) | -0.85 | 0.02 | -45.54 | 0.00 | *** |
|  | Garden: IFG | -0.36 | 0.03 | -13.58 | 0.00 | *** |
| **Component** | **Term** |  |  |  |  |  |
| B. smooth terms | s(tdiff2) | 4.40 | 5.33 | 11.42 | 0.00 | *** |
|  | Smooth term (2014 performance) | 2.44 | 3.08 | 9.13 | 0.00 | *** |
|  | Smooth term (Family) | 0.00 | 621.00 | 0.00 | 0.59 |  |
|  | Smooth term (Locality) | 44.46 | 94.00 | 0.97 | 0.00 | *** |
| Signif. codes: 0 <= '***' < 0.001 < '**' < 0.01 < '*' < 0.05 | | | | | | |
| Adjusted R-squared: 0.238, Deviance explained 0.267 | | | | | | |
| fREML : -66.802, Scale est: 0.0481, N: 1229 | | | | | | |

Table S8: Full model summary for GAM measuring interannual variation in effects of mean winter temperature transfer distance on individual relative growth rates over time. Planting block, nested within garden, was included to account for unintended environmental differences within gardens but is not shown here for easier interpretation.

| **Component** | **Term** | **Estimate** | **Std Error** | **t-value** | **p-value** |  |
| --- | --- | --- | --- | --- | --- | --- |
| A. parametric coefficients | (Intercept) | -1.39 | 0.03 | -44.20 | 0.00 | *** |
|  | Year2016 | 0.76 | 0.03 | 26.92 | 0.00 | *** |
|  | Year2017 | -0.05 | 0.04 | -1.29 | 0.20 |  |
|  | Year2019 | 0.04 | 0.04 | 1.21 | 0.23 |  |
|  | Year2020 | 0.10 | 0.03 | 3.23 | 0.00 | ** |
|  | Year2021 | -0.29 | 0.03 | -8.39 | 0.00 | *** |
|  | Year2022 | 0.23 | 0.03 | 7.23 | 0.00 | *** |
|  | Year2023 | -0.12 | 0.04 | -3.25 | 0.00 | ** |
|  | Year2024 | 0.17 | 0.03 | 5.09 | 0.00 | *** |
|  | SiteBlockChico2 | -0.16 | 0.02 | -6.58 | 0.00 | *** |
|  | SiteBlockChico3 | -0.24 | 0.02 | -10.85 | 0.00 | *** |
|  | SiteBlockChico4 | -0.30 | 0.02 | -14.18 | 0.00 | *** |
|  | SiteBlockChico5 | -0.24 | 0.02 | -11.72 | 0.00 | *** |
|  | SiteBlockIFG1 | -0.42 | 0.02 | -17.92 | 0.00 | *** |
|  | SiteBlockIFG2 | -0.44 | 0.02 | -18.45 | 0.00 | *** |
|  | SiteBlockIFG3 | -0.48 | 0.03 | -18.84 | 0.00 | *** |
|  | SiteBlockIFG4 | -0.42 | 0.02 | -17.78 | 0.00 | *** |
|  | SiteBlockIFG5 | -0.51 | 0.02 | -20.51 | 0.00 | *** |
| **Component** | **Term** |  |  |  |  |  |
| B. smooth terms | s(tdiffyr_win):Year2015 | 1.05 | 1.09 | 0.43 | 0.49 |  |
|  | s(tdiffyr_win):Year2016 | 6.20 | 7.09 | 9.76 | 0.00 | *** |
|  | s(tdiffyr_win):Year2017 | 3.21 | 3.91 | 7.84 | 0.00 | *** |
|  | s(tdiffyr_win):Year2019 | 5.54 | 6.44 | 20.20 | 0.00 | *** |
|  | s(tdiffyr_win):Year2020 | 6.82 | 7.68 | 29.80 | 0.00 | *** |
|  | s(tdiffyr_win):Year2021 | 5.77 | 6.74 | 23.06 | 0.00 | *** |
|  | s(tdiffyr_win):Year2022 | 2.88 | 3.57 | 8.43 | 0.00 | *** |
|  | s(tdiffyr_win):Year2023 | 3.21 | 3.89 | 18.31 | 0.00 | *** |
|  | s(tdiffyr_win):Year2024 | 7.45 | 8.19 | 26.76 | 0.00 | *** |
|  | Smooth term (Year-1 height) | 12.65 | 13.59 | 429.67 | 0.00 | *** |
|  | Smooth term (2014 height) | 3.00 | 3.76 | 4.63 | 0.00 | ** |
|  | Smooth term (Family) | 96.18 | 626.00 | 0.19 | 0.00 | ** |
|  | Smooth term (Locality) | 70.05 | 94.00 | 2.85 | 0.00 | *** |
| Signif. codes: 0 <= '***' < 0.001 < '**' < 0.01 < '*' < 0.05 | | | | | | |
| Adjusted R-squared: 0.585, Deviance explained 0.556 | | | | | | |
| fREML : 23462.302, Scale est: 0.443, N: 24156 | | | | | | |

Table S9: Full model summary for GAM measuring interannual variation in effects of annual maximum temperature transfer distance on individual relative growth rates over time.

| **Component** | **Term** | **Estimate** | **Std Error** | **t-value** | **p-value** |  |
| --- | --- | --- | --- | --- | --- | --- |
| A. parametric coefficients | (Intercept) | -1.35 | 0.02 | -56.97 | 0.00 | *** |
|  | Year2016 | 0.69 | 0.02 | 32.11 | 0.00 | *** |
|  | Year2017 | -0.04 | 0.03 | -1.58 | 0.11 |  |
|  | Year2019 | 0.02 | 0.03 | 0.82 | 0.41 |  |
|  | Year2020 | -0.01 | 0.03 | -0.33 | 0.74 |  |
|  | Year2021 | -0.32 | 0.03 | -9.58 | 0.00 | *** |
|  | Year2022 | 0.13 | 0.03 | 4.49 | 0.00 | *** |
|  | Year2023 | -0.12 | 0.03 | -3.80 | 0.00 | *** |
|  | Year2024 | 0.16 | 0.04 | 4.52 | 0.00 | *** |
|  | SiteBlockChico2 | -0.14 | 0.02 | -5.53 | 0.00 | *** |
|  | SiteBlockChico3 | -0.23 | 0.02 | -10.31 | 0.00 | *** |
|  | SiteBlockChico4 | -0.29 | 0.02 | -13.46 | 0.00 | *** |
|  | SiteBlockChico5 | -0.23 | 0.02 | -11.46 | 0.00 | *** |
|  | SiteBlockIFG1 | -0.32 | 0.02 | -13.82 | 0.00 | *** |
|  | SiteBlockIFG2 | -0.35 | 0.02 | -14.80 | 0.00 | *** |
|  | SiteBlockIFG3 | -0.38 | 0.03 | -15.16 | 0.00 | *** |
|  | SiteBlockIFG4 | -0.33 | 0.02 | -14.12 | 0.00 | *** |
|  | SiteBlockIFG5 | -0.41 | 0.02 | -16.55 | 0.00 | *** |
| **Component** | **Term** |  |  |  |  |  |
| B. smooth terms | s(tdiffyr):Year2015 | 5.48 | 6.46 | 39.01 | 0.00 | *** |
|  | s(tdiffyr):Year2016 | 6.68 | 7.67 | 13.17 | 0.00 | *** |
|  | s(tdiffyr):Year2017 | 2.14 | 2.68 | 7.46 | 0.00 | *** |
|  | s(tdiffyr):Year2019 | 3.71 | 4.59 | 12.28 | 0.00 | *** |
|  | s(tdiffyr):Year2020 | 6.27 | 7.34 | 11.68 | 0.00 | *** |
|  | s(tdiffyr):Year2021 | 4.16 | 5.01 | 29.36 | 0.00 | *** |
|  | s(tdiffyr):Year2022 | 1.00 | 1.00 | 3.54 | 0.06 | . |
|  | s(tdiffyr):Year2023 | 1.68 | 2.09 | 13.07 | 0.00 | *** |
|  | s(tdiffyr):Year2024 | 3.23 | 3.90 | 39.09 | 0.00 | *** |
|  | Smooth term (Year-1 height) | 12.48 | 13.49 | 423.13 | 0.00 | *** |
|  | Smooth term (2014 height) | 3.00 | 3.76 | 4.33 | 0.00 | ** |
|  | Smooth term (Family) | 97.68 | 626.00 | 0.19 | 0.00 | ** |
|  | Smooth term (Locality) | 49.70 | 94.00 | 1.43 | 0.00 | *** |
| Signif. codes: 0 <= '***' < 0.001 < '**' < 0.01 < '*' < 0.05 | | | | | | |
| Adjusted R-squared: 0.588, Deviance explained 0.557 | | | | | | |
| fREML : 23389.286, Scale est: 0.444, N: 24156 | | | | | | |

Table S10: Full model summary for GAM measuring interannual variation in effects of averaged summer maximum temperature transfer distance on individual relative growth rates over time.

| **Component** | **Term** | **Estimate** | **Std Error** | **t-value** | **p-value** |  |
| --- | --- | --- | --- | --- | --- | --- |
| A. parametric coefficients | (Intercept) | -1.43 | 0.02 | -65.33 | 0.00 | *** |
|  | Year2016 | 0.74 | 0.02 | 38.83 | 0.00 | *** |
|  | Year2017 | 0.04 | 0.02 | 1.59 | 0.11 |  |
|  | Year2019 | 0.12 | 0.03 | 4.75 | 0.00 | *** |
|  | Year2020 | 0.06 | 0.03 | 2.46 | 0.01 | * |
|  | Year2021 | -0.20 | 0.03 | -5.83 | 0.00 | *** |
|  | Year2022 | 0.21 | 0.03 | 8.00 | 0.00 | *** |
|  | Year2023 | -0.08 | 0.03 | -2.77 | 0.01 | ** |
|  | Year2024 | 0.11 | 0.03 | 4.02 | 0.00 | *** |
|  | SiteBlockChico2 | -0.13 | 0.02 | -5.35 | 0.00 | *** |
|  | SiteBlockChico3 | -0.22 | 0.02 | -10.76 | 0.00 | *** |
|  | SiteBlockChico4 | -0.28 | 0.02 | -13.14 | 0.00 | *** |
|  | SiteBlockChico5 | -0.22 | 0.02 | -10.99 | 0.00 | *** |
|  | SiteBlockIFG1 | -0.35 | 0.02 | -14.89 | 0.00 | *** |
|  | SiteBlockIFG2 | -0.38 | 0.02 | -15.71 | 0.00 | *** |
|  | SiteBlockIFG3 | -0.42 | 0.03 | -16.42 | 0.00 | *** |
|  | SiteBlockIFG4 | -0.37 | 0.02 | -15.27 | 0.00 | *** |
|  | SiteBlockIFG5 | -0.44 | 0.02 | -17.52 | 0.00 | *** |
| **Component** | **Term** |  |  |  |  |  |
| B. smooth terms | s(tdiffyr):Year2015 | 5.72 | 6.74 | 33.12 | 0.00 | *** |
|  | s(tdiffyr):Year2016 | 6.37 | 7.39 | 11.13 | 0.00 | *** |
|  | s(tdiffyr):Year2017 | 2.33 | 2.92 | 10.52 | 0.00 | *** |
|  | s(tdiffyr):Year2019 | 4.86 | 5.87 | 15.55 | 0.00 | *** |
|  | s(tdiffyr):Year2020 | 5.88 | 6.91 | 19.52 | 0.00 | *** |
|  | s(tdiffyr):Year2021 | 4.06 | 4.86 | 36.11 | 0.00 | *** |
|  | s(tdiffyr):Year2022 | 1.00 | 1.00 | 10.54 | 0.00 | ** |
|  | s(tdiffyr):Year2023 | 1.99 | 2.50 | 22.25 | 0.00 | *** |
|  | s(tdiffyr):Year2024 | 3.05 | 3.78 | 50.40 | 0.00 | *** |
|  | Smooth term (Year-1 height) | 12.45 | 13.49 | 465.53 | 0.00 | *** |
|  | Smooth term (Family) | 102.04 | 626.00 | 0.20 | 0.00 | *** |
|  | Smooth term (Locality) | 50.25 | 94.00 | 1.51 | 0.00 | *** |
| Signif. codes: 0 <= '***' < 0.001 < '**' < 0.01 < '*' < 0.05 | | | | | | |
| Adjusted R-squared: 0.593, Deviance explained 0.560 | | | | | | |
| fREML : 23228.617, Scale est: 0.439, N: 24147 | | | | | | |

Table S11: Full model summary of repeated measures ANOVA on yearly relative growth rate by family (cooler garden).

| stratum | term | df | sumsq | meansq | statistic | p.value |
| --- | --- | --- | --- | --- | --- | --- |
| character | character | numeric | numeric | numeric | numeric | numeric |
| Accession_progeny | Accession | 621 | 32.5 | 0.1 | 2.8 | 0.0 |
| Accession_progeny | Year | 9 | 3.0 | 0.3 | 18.1 | 0.0 |
| Accession_progeny | SiteBlock | 4 | 7.1 | 1.8 | 96.4 | 0.0 |
| Accession_progeny | Accession:Year | 982 | 44.6 | 0.0 | 2.5 | 0.0 |
| Accession_progeny | Residuals | 39 | 0.7 | 0.0 |  |  |
| Within | Year | 9 | 246.5 | 27.4 | 473.3 | 0.0 |
| Within | Accession:Year | 5,329 | 307.2 | 0.1 | 1.0 | 0.6 |
| Within | Residuals | 5,900 | 341.5 | 0.1 |  |  |
| n: 8 | | | | | | |

Table S12: Full model summary of repeated measures ANOVA on yearly relative growth rate by family (warmer garden).

| stratum | term | df | sumsq | meansq | statistic | p.value |
| --- | --- | --- | --- | --- | --- | --- |
| character | character | numeric | numeric | numeric | numeric | numeric |
| Accession_progeny | Accession | 624 | 48.8 | 0.1 | 2.0 | 0.0 |
| Accession_progeny | Year | 9 | 9.3 | 1.0 | 27.2 | 0.0 |
| Accession_progeny | SiteBlock | 4 | 13.6 | 3.4 | 89.1 | 0.0 |
| Accession_progeny | Accession:Year | 1,038 | 50.1 | 0.0 | 1.3 | 0.2 |
| Accession_progeny | Residuals | 42 | 1.6 | 0.0 |  |  |
| Within | Year | 9 | 1,238.2 | 137.6 | 1,364.7 | 0.0 |
| Within | Accession:Year | 5,354 | 432.9 | 0.1 | 0.8 | 1.0 |
| Within | Residuals | 6,345 | 639.6 | 0.1 |  |  |
| n: 8 | | | | | | |

Table S13: Full model summary of ANOVA and Tukey post-hoc comparison of seed sourcing outcomes modeled via simulated transfer of families from the cooler garden to the warmer garden.

| group |  | | Estimate | Standard Error | |  | | statistic | |  |  |  |
| --- | --- | --- | --- | --- | --- | --- | --- | --- | --- | --- | --- | --- |
| Fixed effects | | | | | | | | | |  |  |  |
|  | (Intercept) | | 0.404 | 0.015 | |  | | 26.534 | |  |  |  |
|  | approach0.5 | | 0.015 | 0.020 | |  | | 0.732 | |  |  |  |
|  | approach1 | | 0.070 | 0.029 | |  | | 2.403 | |  |  |  |
|  | approach3 | | 0.062 | 0.022 | |  | | 2.807 | |  |  |  |
| Random effects | | | | | | | | | |  |  |  |
| Locality | sd__(Intercept) | | 0.045 |  | |  | |  | |  |  |  |
| Residual | sd__Observation | | 0.140 |  | |  | |  | |  |  |  |
| square root of the estimated residual variance: 0.1 | | | | | | | | | |  |  |  |
| data's log-likelihood under the model: 175.5 | | | | | | | | | |  |  |  |
| Akaike Information Criterion: -338.9 | | | | | | | | | |  |  |  |
| Bayesian Information Criterion: -315.5 | | | | | | | | | |  |  |  |
| term | | contrast | | | null.value | | estimate | | std.error | | statistic | adj.p.value |
| character | | character | | | numeric | | numeric | | numeric | | numeric | numeric |
| approach | | Climate match - random | | | 0 | | 0.0 | | 0.0 | | 0.7 | 0.9 |
| approach | |  | | | 0 | | 0.1 | | 0.0 | | 2.4 | 0.1 |
| approach | | Phenotypic selection - random | | | 0 | | 0.1 | | 0.0 | | 2.8 | 0.0 |
| approach | |  | | | 0 | | 0.1 | | 0.0 | | 1.9 | 0.2 |
| approach | | Phenotypic selection - climate match | | | 0 | | 0.0 | | 0.0 | | 2.1 | 0.1 |
| approach | |  | | | 0 | | -0.0 | | 0.0 | | -0.3 | 1.0 |
| n: 6 | | | | | | | | | | | | |

Table S14: Cohorts (A-E) of individuals in two common gardens based on temperature difference ranges (negative temperature difference: trees are from warmer environments than the garden, zero temperature difference: trees are from environments with equal temperature to the garden, positive temperature difference: trees are from cooler environments than the garden). Cohorts are defined differently for each garden (C subscript: cooler garden, W subscript: warmer garden) based on the distribution of temperature differences (Fig. S4), resulting in temperature ranges that are not necessarily equal or contiguous between cohorts.

| Garden | Cohort | N | Temperature difference range (°C) |
| --- | --- | --- | --- |
| character | character | integer | character |
| Cooler (IFG) | AC | 64 | (-4.67, -2] |
|  | BC | 246 | (-2, 0] |
|  | CC | 145 | (0, 2] |
|  | DC | 139 | (2, 5] |
|  | EC | 59 | (5, 8.9] |
| Warmer (CSO) | AW | 31 | (-2.05, 0] |
|  | BW | 201 | (0, 2] |
|  | CW | 105 | (3, 5] |
|  | DW | 109 | (5, 7] |
|  | EW | 69 | (7, 9.08] |
| n: 10 | | | |

Table S15: Full model summary of repeated measures ANOVA on yearly performance by cohort (cooler garden).

| stratum | term | df | sumsq | meansq | statistic | p.value |
| --- | --- | --- | --- | --- | --- | --- |
| character | character | numeric | numeric | numeric | numeric | numeric |
| Accession | cohort | 4 | 7.9 | 2.0 | 11.8 | 0.0 |
| Accession | Year | 2 | 1.2 | 0.6 | 3.7 | 0.0 |
| Accession | Residuals | 643 | 107.1 | 0.2 |  |  |
| Within | Year | 9 | 1.9 | 0.2 | 53.4 | 0.0 |
| Within | cohort:Year | 36 | 1.4 | 0.0 | 9.6 | 0.0 |
| Within | Residuals | 5,803 | 23.0 | 0.0 |  |  |
| n: 6 | | | | | | |

Table S16: Full model summary of repeated measures ANOVA on yearly performance by cohort (warmer garden).

| stratum | term | df | sumsq | meansq | statistic | p.value |
| --- | --- | --- | --- | --- | --- | --- |
| character | character | numeric | numeric | numeric | numeric | numeric |
| Accession | cohort | 4 | 2.0 | 0.5 | 3.5 | 0.0 |
| Accession | Year | 2 | 2.3 | 1.2 | 7.9 | 0.0 |
| Accession | cohort:Year | 1 | 0.0 | 0.0 | 0.0 | 0.9 |
| Accession | Residuals | 501 | 73.2 | 0.1 |  |  |
| Within | Year | 7 | 1.6 | 0.2 | 70.7 | 0.0 |
| Within | cohort:Year | 28 | 0.4 | 0.0 | 4.2 | 0.0 |
| Within | Residuals | 3,519 | 11.7 | 0.0 |  |  |
| n: 7 | | | | | | |

Table S17: Full model summary of geographic and climatic effects on cumulative performance in 2024 at the cooler garden.

|  | Estimate | Standard Error | t value | Pr(>\|t\|) |  |
| --- | --- | --- | --- | --- | --- |
| (Intercept) | 0.303 | 0.006 | 50.089 | 0.0000 | *** |
| scale(PC1) | 0.044 | 0.007 | 6.169 | 0.0000 | *** |
| scale(PC2) | -0.026 | 0.009 | -2.890 | 0.0040 | ** |
| scale(Latitude) | -0.003 | 0.015 | -0.189 | 0.8499 |  |
| scale(Longitude) | -0.022 | 0.015 | -1.474 | 0.1409 |  |
| *Signif. codes: 0 <= '***' < 0.001 < '**' < 0.01 < '*' < 0.05* | | | | | |
| Residual standard error: 0.1543 on 645 degrees of freedom | | | | | |
| Multiple R-squared: 0.08144, Adjusted R-squared: 0.07574 | | | | | |
| F-statistic: 14.3 on 645 and 4 DF, p-value: 0.0000 | | | | | |

Table S18: Full model summary of geographic and climatic effects on cumulative performance in 2024 at the warmer garden.

|  | Estimate | Standard Error | t value | Pr(>\|t\|) |  |
| --- | --- | --- | --- | --- | --- |
| (Intercept) | 0.403 | 0.006 | 68.658 | 0.0000 | *** |
| scale(PC1) | 0.020 | 0.007 | 2.884 | 0.0041 | ** |
| scale(PC2) | -0.024 | 0.009 | -2.819 | 0.0050 | ** |
| scale(Latitude) | -0.001 | 0.015 | -0.094 | 0.9251 |  |
| scale(Longitude) | -0.018 | 0.014 | -1.227 | 0.2201 |  |
| *Signif. codes: 0 <= '***' < 0.001 < '**' < 0.01 < '*' < 0.05* | | | | | |
| Residual standard error: 0.1471 on 624 degrees of freedom | | | | | |
| Multiple R-squared: 0.04191, Adjusted R-squared: 0.03577 | | | | | |
| F-statistic: 6.824 on 624 and 4 DF, p-value: 0.0000 | | | | | |

Table S19: Full model summary of geographic and climatic effects on cumulative relative growth rate across the study period at the cooler garden.

|  | Estimate | Standard Error | t value | Pr(>\|t\|) |  |
| --- | --- | --- | --- | --- | --- |
| (Intercept) | 0.175 | 0.002 | 96.835 | 0.0000 | *** |
| scale(PC1) | 0.002 | 0.002 | 0.957 | 0.3387 |  |
| scale(PC2) | -0.003 | 0.003 | -1.232 | 0.2181 |  |
| scale(Latitude) | 0.019 | 0.005 | 4.144 | 0.0000 | *** |
| scale(Longitude) | 0.014 | 0.004 | 3.227 | 0.0013 | ** |
| *Signif. codes: 0 <= '***' < 0.001 < '**' < 0.01 < '*' < 0.05* | | | | | |
| Residual standard error: 0.07374 on 1662 degrees of freedom | | | | | |
| Multiple R-squared: 0.03604, Adjusted R-squared: 0.03372 | | | | | |
| F-statistic: 15.53 on 1662 and 4 DF, p-value: 0.0000 | | | | | |

Table S20: Full model summary of geographic and climatic effects on cumulative relative growth rate across the study period at the warmer garden.

|  | Estimate | Standard Error | t value | Pr(>\|t\|) |  |
| --- | --- | --- | --- | --- | --- |
| (Intercept) | 0.219 | 0.002 | 128.344 | 0.0000 | *** |
| scale(PC1) | -0.005 | 0.002 | -2.621 | 0.0089 | ** |
| scale(PC2) | 0.000 | 0.003 | 0.033 | 0.9733 |  |
| scale(Latitude) | 0.013 | 0.004 | 2.925 | 0.0035 | ** |
| scale(Longitude) | 0.009 | 0.004 | 2.115 | 0.0346 | * |
| *Signif. codes: 0 <= '***' < 0.001 < '**' < 0.01 < '*' < 0.05* | | | | | |
| Residual standard error: 0.07095 on 1719 degrees of freedom | | | | | |
| Multiple R-squared: 0.01583, Adjusted R-squared: 0.01354 | | | | | |
| F-statistic: 6.913 on 1719 and 4 DF, p-value: 0.0000 | | | | | |
